# JNK signaling regulates reproductive trade-offs after *Plasmodium* infection in the malaria mosquito

**DOI:** 10.1101/2025.09.03.673998

**Authors:** Max Lombardi, Anastasia Accotti, Paolo Scarpelli, Gloria Iacomelli, Lucia Pazzagli, Antonella Turco, Giulia Monacchia, Simona Ferracchiato, Tasneem A. Rinvee, Duo Peng, Charles Vidoudez, W. Robert Shaw, Flaminia Catteruccia, Roberta Spaccapelo, Matthew J. Peirce

## Abstract

Environmental stress can limit mammalian reproduction by affecting production of sexual steroid hormones. Here we reveal a similar mechanism in the malarial mosquito *Anopheles gambiae*: activation of the stress-sensitive c-<u>J</u>un <u>N</u>-terminal <u>k</u>inase, JNK, constrains reproductive investment by suppressing production of ecdysteroids that orchestrate egg development in this species. We show that infection with *Plasmodium berghei* parasites increases JNK signalling in the reproductive tract causing a JNK-dependent reduction in both egg development and mosquito survival. Moreover, JNK signaling supresses expression of *Cyp315a1* (AGAP000284), a rate-limiting enzyme in ecdysteroid synthesis, a transcriptional change reflected in reduced ecdysteroid production following an infected blood meal. A similar mechanism limits egg production under other stressors (heat stress, or ectopic activation of JNK signaling). Together, these data reveal a regulatory circuit whereby *Plasmodium* infection curtails reproductive investment in an important vector of human malaria, one that may be applicable to environmental stressors more generally.

## INTRODUCTION

Exposure to environmental stresses can modify investment in reproduction across taxa [1–4]. In the particular case of female anopheline mosquitoes, such trade-offs between stress resilience and reproduction impact directly on their capacity to act as a vector for *Plasmodium* parasites which cause malaria, a disease that continues to impose an intolerable global public health burden [5]. Investment in immunity and stress resistance underpins mosquito longevity and susceptibility to infection that are central to the likelihood of parasite transmission. At the same time, investment in reproductive output controls population size and thus transmission frequency. Despite their importance to vector capacity, understanding of the mechanisms and signaling pathways that regulate these trade-offs remains limited.

*Anopheles gambiae* females invest in reproduction through the production of lipid- and protein-rich eggs, a costly endeavor sustained by the nutritional resources contained in a blood meal. The pathways overseeing this investment in egg development comprise several components [6]. Insulin-like peptides, produced in the brain in response to blood feeding [7–9], act systemically though a single insulin-like receptor [10]. In addition, free amino acids, liberated by tryptic digestion of the blood meal, act in combination with insulin to activate growth signals through the Target of Rapamycin (ToR) pathway [11]. Meanwhile, these nutritionally-derived blood feeding signals are complemented by the production of ovarian ecdysteroid hormone (OEH) [12], which acts in concert with insulin-like peptides [13] to stimulate the synthesis and release of ecdysone (E) [14] from ovarian epithelial cells. E is then hydroxylated at the C20 position in the fat body to produce the active hormone 20-hydroxyecdysone (20E), which acts through its heterodimeric receptor comprising ecdysone receptor (EcR) and ultraspiracle (USP) [15] to drive egg development. Disabling 20E production, or either component of its receptor, profoundly inhibits egg production [16].

When the blood meal is infected with *Plasmodium* parasites, the resources contained within it must be shared between supporting egg development and raising an energetically costly anti-parasitic immune response. Indeed, fitness costs associated with infection are well documented; a *Plasmodium-*infected blood meal frequently leads to a reduction in the number of eggs developed [17,18] as well as an increase in female mortality [19]. While the pathways mediating these costs of *Plasmodium* infection remain only partially elucidated, some mechanistic clues are provided by particular parasite/ mosquito combinations, in which fitness costs of infection are reduced or absent. In ‘natural’ pairings—those in which parasite and mosquito host share extensive evolutionary history—reproductive costs associated with infection appear to be minimal [19]. A case in point is that of *Plasmodium falciparum* and *Anopheles gambiae* in which infection has no effect on mosquito fecundity [20]. This apparently mutualistic relationship may be linked to the *Plasmodium falciparum* surface protein of 47kDa (Pfs47) which is highly strain-specific [21] and allows the parasite, in the context of its natural mosquito host, to avoid provoking a mosquito immune response by silencing the immune- and stress-activated kinase, JNK (c-Jun N-terminal Kinase) [22,23]. In contrast, *P. falciparum* infection of non-adapted anophelines or infection of *An. gambiae* with non-adapted *Plasmodium* strains [22], such as the rodent malaria parasite *P. berghei*, result in their JNK-dependent destruction [24]. Similar to reproductive costs, the infection-induced increase in mosquito mortality, is also confined to non-natural infections [19].

Beyond *Plasmodium* infection, mitigating against other immune challenge or environmental stresses may also cause competition for the resources derived from a blood meal. Bacterial infection, non-infectious immune challenge, as well as heat- or cold-stress have all been shown to impact fecundity [17,25]. Moreover, some mosquitoes, including anophelines [26], can undergo complete reproductive arrest (diapause) in order to conserve resources to favor survival during dry or cold periods [27]. Thus, mechanisms that allow mosquitoes to redistribute resources between reproduction and stress resilience clearly exist, even if the molecular details remain incomplete.

In this context, JNK may be of interest. JNK is part of an ancient, stress-activated signaling cascade, sensitive to an array of stressful environmental stimuli including trauma [28] inflammation [29], heat [30,31] and oxidative stress [32]. JNK phosphorylates downstream transcription factors such as Jun, Fos [33] and FoxO [34,35], activating their transcriptional activity and driving the expression of an array of genes linked to wound healing [36], heat-shock [31,32,37] and redox regulation [32] that mitigate the impact of the initiating stress, promoting survival and longevity [32,35]. At least one target of the JNK pathway, *puckered*, constitutes a negative feedback loop. By acting as a JNK phosphatase, Puckered dephosphorylates JNK, returning its activity to baseline [38].

While promoting stress resilience, JNK supresses investment in growth across genera. JNK activation inhibits insulin-mediated growth and metabolic reprograming in *Drosophila* [35] and *C. elegans* [37] while in mammals elevated JNK activity is causally linked to the insulin insensitivity and consequent ‘metabolic syndrome’ that characterize type 2 diabetes [39,40]. Interestingly, in polycystic ovarian syndrome, a condition also linked to obesity and insulin insensitivity [41], JNK is implicated in the dysregulation of steroid sex hormones that underpins the infertility characteristic of the disease [42]. JNK signaling antagonizes insulin at multiple levels, inhibiting its production in the insect brain [35,37], its signaling through the inhibitory phosphorylation of insulin receptor substrate 1 (IRS1) [43] as well as its transcriptional effects by activating the insulin inhibitory transcription factor, FoxO [34,37].

Here we identify an important role for the JNK pathway in regulating the production of ecdysteroids to control the division of resources between reproduction and stress mitigation in the major malaria vector *Anopheles gambiae*, with significant implications for its capacity to transmit malaria-causing parasites.

## RESULTS

### Plasmodium berghei infection induces fitness costs

To investigate the molecular mechanisms underpinning fitness costs caused by *Plasmodium* infection, we chose a non-natural model system in which such costs have previously been reported [17,18,44]. In line with these studies [18,44], using the G3 strain of *An. gambiae* (a natural vector for *P. falciparum*) and the rodent malaria parasite *P. berghei,* we observed that virgin females blood feeding on a *P. berghei*-infected mouse developed significantly fewer eggs (**Supp. Fig. 1A**) and were also significantly more likely to die in the first 8 days following infection (**Supp. Fig. 1B**) compared to mosquitoes fed on an uninfected littermate control. In mated females, *P. berghei* infection also reduced the number of eggs oviposited (**Supp. Fig. 1C**). Consistent with previous reports [18,44], mosquitoes subjected to an infected blood meal, then maintained at a temperature non-permissive to *P. berghei* growth (27°C), exhibited no such reduction in fecundity suggesting that the reduced egg development observed was attributable to the development of the parasite rather than any reduction in nutritional value of the blood meal or to the murine anti-parasite immune response (**Supp. Fig. 1D**).

### P. berghei-induced JNK signaling is linked to fitness costs of infection

These fitness costs associated with *P. berghei* infection of *An. gambiae* are not observed with *P. falciparum* [16,20]. One difference between the two parasites lies in their respective abilities to activate the JNK pathway during infection [23,24]. We hypothesized that any role for JNK in the reduction in egg numbers following *P. berghei* infection might be reflected in increased JNK signaling in tissues involved in egg development. To address this possibility, we examined the presence of active (phosphorylated) JNK (pJNK) in the reproductive tracts (RT) (comprising ovaries, atrium and spermatheca) of unfed females or those fed on infected or uninfected mice. At 48 hours post blood feeding (hPBF) we observed an increase in pJNK levels in the RT of females fed on infected blood, (**Figs. 1A, 1B**) that was statistically significant using a one sample t test (*p*=0.0038, t=8.2, df=3 discrepancy=0.598, 95% CI=0.37-0.83). In contrast, levels of pS6K, a kinase at which growth signals downstream of insulin and ToR-like signaling pathways are integrated [45], were increased by blood feeding as expected [46] but unaffected by the presence of *P. berghei* (*p*=0.64, t=0.53, df=3, discrepancy=0.12, 95% CI=-0.85-0.61)

**Fig 1.**
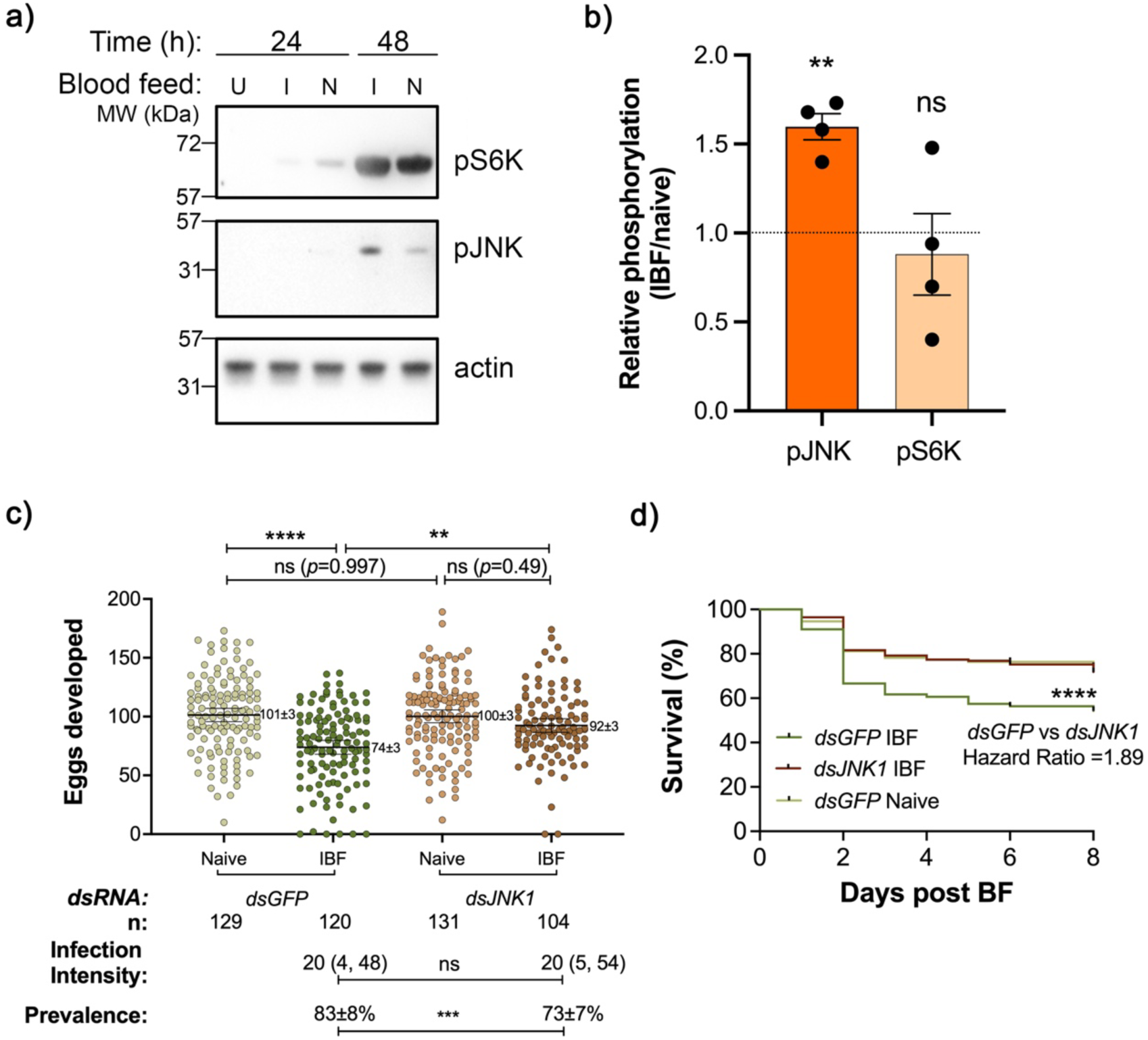
Increased *JNK* activation with *P. berghei* linked to fitness costs of infection. **a)** A representative western blot for pJNK, pS6K and actin in reproductive tracts dissected 24 or 48 hPBF on a *P. berghei*-infected (I), uninfected naïve mouse (N) or from unfed controls (U). Originals of the western blots presented here appear in full in Supplemantary Data**. b)** Actin-normalized densitometry values for pJNK and pS6K were used to calculate the ratio of the signal in infected vs uninfected blood-fed females. Deviation from a ratio of 1 (indicated by the dashed line) was compared using a one sample t-test (test statistics given in text). Data show the mean ± SEM of 4 independent biological replicates, each represented by a black circle. **c)** Virgin females were injected with the indicated *dsRNA* and allowed to feed on a *P. berghei*-infected (IBF) or an uninfected (Naïve) mouse. At day 8 PBF, eggs and oocysts were counted. Uninfected females (those with eggs but no oocysts) were excluded from the egg counts. Data represent the mean ± 95% confidence interval of 6 biological replicates. The mean ± SEM is also shown numerically. The total number of mosquitoes analyzed (n), median infection intensity and interquartile range and mean ± SEM % prevalence are reported below the graph. Differences in egg number were compared using a negative binomial generalized linear model with Tukey’s correction of pairwise comparisons (see Methods). Selected comparisons are indicated: *dsGFP*-IBF/*dsJNK1-*IBF(effect ratio=0.80, 95%CI [0.69, 0.93], Z=-3.84, *p*=0.00071); *dsGFP*-IBF/*dsGFP*-naive (effect ratio=0.73, 95%CI [0.63, 0.84], Z= -5.73, *p*<0.0001); d*sJNK1*-IBF/*dsJNK1*-naive (effect ratio=0.92, 95%CI [0.49-0.95, Z=-1.42, *p*=0.48]. Differences in infection intensity and prevalence were respectively compared using a Mann-Whitney test and a Fisher’s exact test on the absolute prevalence data. **d)** Daily survival rates from 4 independent biological replicates were compared using individual Log-rank (Mantel-Cox) tests. Survival was significantly different in the *dsJNK1*-IBF and *dsGFP*-IBF groups (as indicated, p<0.0001, Chi square= 34.26, df=1, HR=1.89, 95% CI [1.51, 2.37]) and *dsGFP*-naïve and *dsGFP*-IBF groups (p<0.0001, Chi square= 34.14, HR=1.97, 95% CI [1.56, 2.48] ) but indistinguishable in the *dsGFP*-naïve and *dsJNK1*-IBF groups (*p*=0.75). **** indicates *p*<0.0001; **, *p*<0.01; ns, ‘not significant’ (*p*>0.05).

To examine the possibility of a causal link between increased JNK signaling in the RT and reduced egg development we injected virgin females with ds*JNK1 (AGAP009461)* or *dsGFP* [47], not expressed in these mosquitoes, as a control. *P. berghei* infection in *dsGFP*-injected females resulted in a reduced number of eggs (**Fig. 1C**) and increased mortality PBF (**Fig. 1D**), similar to our results in uninjected females (**Supp. Figs. 1A, 1B**). While *dsJNK1* successfully reduced *JNK1* mRNA expression (**Supp. Fig. 2A**), it had no impact on the number of eggs developed in the absence of infection (**Fig. 1C**) [48]. However, in the *JNK1*-depleted females, both the decrease in egg number (**Fig. 1C**) and the increase in mortality (**Fig. 1D**) observed in *dsGFP* controls following *Plasmodium* infection were completely reversed. In previous studies [24] the depletion of JNK led to an increase in oocyst numbers that we did not observe in our pooled data (**Fig. 1C**). However, consistent with those reports, in three of six experiments in which our infection intensities and theirs were similar (median 5-10 oocysts/midgut), *JNK1* depletion did indeed increase infection intensity (**Supp. Fig. 3**). We also noted a small but significant reduction in infection prevalence in *JNK1*-depleted females (**Fig. 1C**) that runs counter to published data [24]. Again, this may relate to the higher mean infection intensities in our experiments compared to those published since we noted no *dsJNK1*-induced decrease in prevalence in the three lower intensity infections reported in **Supplementary Figure 3**.

Together, these data suggest that JNK signaling may impact *P. berghei* infection in an intensity-specific manner but also support a causal link between JNK activation during *P. berghei* infection and the resulting fecundity and mortality fitness costs to the mosquito.

### RNA-seq analysis suggests role for JNK1 in immunity and 20E synthesis

To understand how JNK signaling shapes gene expression patterns in the RT in the context of blood feeding, we performed RNA-seq analysis of RT samples from *dsGFP-* and *dsJNK1-*injected females, 24 hours after taking a *P. berghei*-infected or naïve blood meal. This is a time point critical for ookinete invasion of the midgut [49] as well as for transcriptional responses to infection [50]. We first focussed our analysis on those genes differentially expressed in *dsGFP* vs *dsJNK1*-treated females following a *P. berghei*-infected blood meal. As summarised in **Supplementary Figure 4A**, we identified 69 genes whose expression was significantly altered (adjusted *p*<0.05) by *JNK1* depletion in the context of a *P. berghei*-infected blood meal, 52 of which (75.4%) were downregulated and 17 (24.6%) were upregulated in the relative absence of JNK (**Supp. Table 1**). Consistent with the role of JNK in immune and wound healing responses [36,51], 41.4 % of JNK-sensitive genes had a predicted immune function or were enriched in hemocytes, cells akin to mammalian macrophages [52] with important roles in immunity and wound healing [53]. One of these downregulated immune genes was SRPN6, a serine protease inhibitor (serpin) implicated in responses to *P. berghei* infection and defined as a marker of plasmodial midgut invasion [54]. However, we identified no candidate genes with an established role in regulating egg production.

We then performed a second analysis to identify genes differentially expressed in *dsJNK1* vs *dsGFP*-injected females irrespective of whether they had fed on naïve or *Plasmodium*-infected blood. Summarised in **Supplementary Figure 4B**, this yielded a larger group of 118 differentially expressed genes which contained 29 of the 69 genes identified in the IBF-specific analysis, though not SRPN6, and showed a similar bias towards genes linked to immunity and wound healing (43.2%) (**Supp. Table 2**).

### P. berghei infections reduce 20E levels after blood feeding

One of the 89 genes unique to this analysis was the cytochrome p450 enzyme, *Cyp315a1* (*AGAP000284*), whose expression was enhanced by *JNK1* depletion. *Cyp315a1,* also known as *Shadow*, is a member of the *Halloween* family of genes that together catalyze rate-limiting steps in the synthesis of the insect reproductive steroid hormone 20-hydroxyecdysone (20E) from cholesterol [55]. We validated the RNA-seq results by qRT-PCR confirming a significant increase in *Cyp315a1* expression albeit only in those females fed on *P. berghei*-infected blood. We also documented a predicted decrease in two immune-related genes, *TGase2* (*AGAP009098*) and *Eater* (*AGAP012386*), irrespective of the *Plasmodium* status of the blood meal consumed (**Fig. 2A**). Given this finding, we measured ecdysteroid levels to assess the impact of *P. berghei* infection on ecdysteroid production after a blood meal. *P. berghei* infection led to significantly reduced levels of ecdysteroids compared to controls (**Fig. 2B**). Notably, both the decrease in ecdysteroid production and the peak in activation of JNK signaling after blood feeding occurred at 48h PBF (**Fig. 1A**).

**Fig 2.**
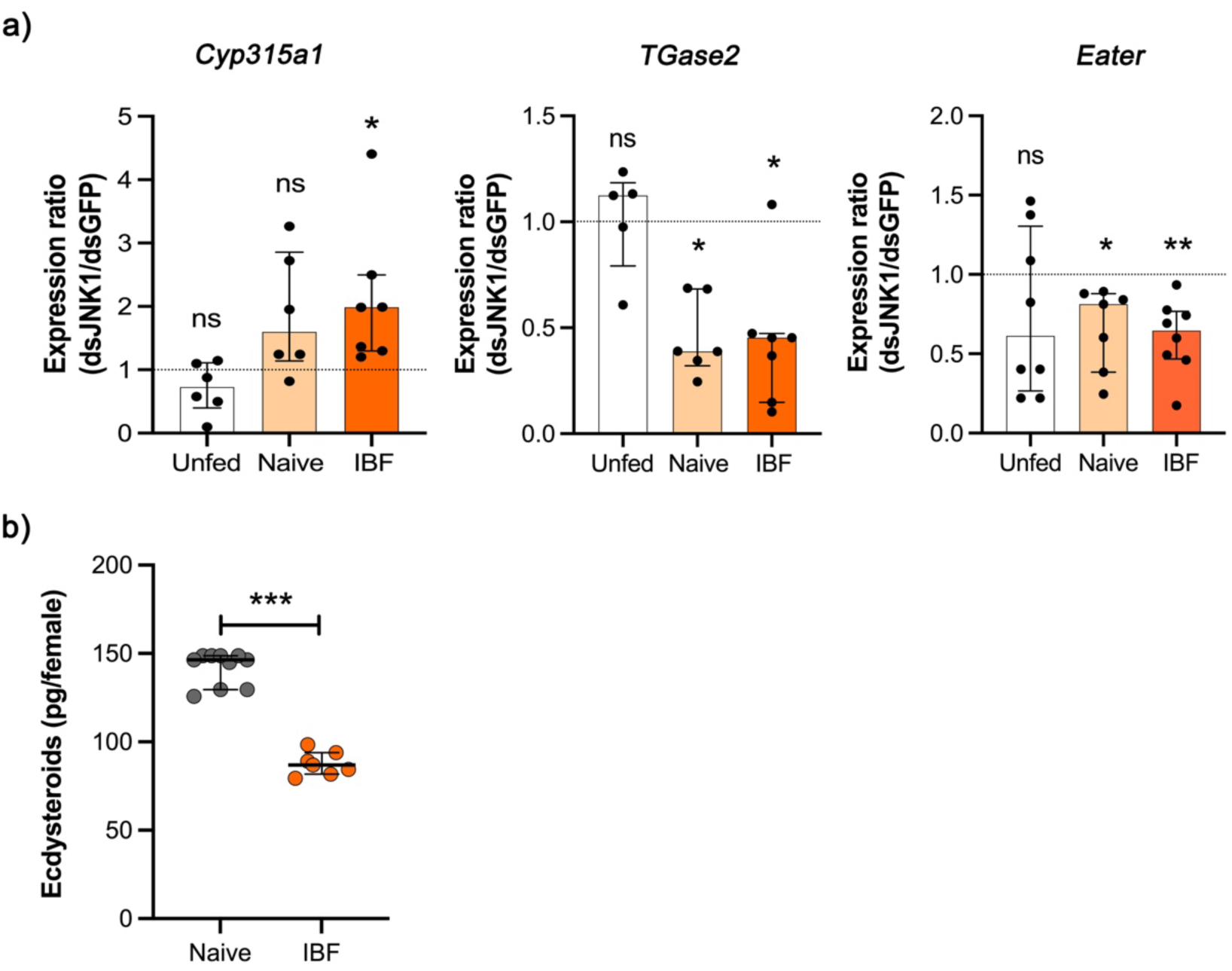
*JNK1*-depleted females invest less in immunity and more in reproduction. **a)** Expression of the three genes indicated was analyzed by qRT-PCR using cDNA prepared from reproductive tracts (ovaries, spermatheca and atrium) dissected from *dsGFP-* or *dsJNK1-*injected females 24 hours after feeding on a *P. berghei*-infected mouse (IBF), an uninfected littermate (Naïve) or controls feeding only on sugar solution (unfed). The relative expression of each gene was used to calculate the ratio of expression in *dsJNK1*-injected vs *dsGFP*-injected females. Deviation from a ratio of 1 (indicated by the dashed line) was assessed using a Wilcoxon signed rank test. *Cyp315a1* (IBF), n=7, *p*=0.016, W=28, discrepancy=0.99, 95%CI [0.20, 3.4]; *TGase2 (*Naïve), n=6, *p*= 0.031, W=-21, discrepancy=-0.61, 95% CI [-0.75, -0.31]) (IBF), n=7, *p*=0.031, W=-26, discrepancy=-0.55, 95%CI [-0.90, - 0.081]; *Eater* (Naïve) n=7, *p*=0.016, discrepancy=-0.19, 95%CI [-0.75, -0.10]; (IBF), n=8, *p*=0.0078, W=-36, discrepancy= -0.36, 95% CI [-0.83, -0.067]. Data shown represent the median ± interquartile range of at least 5 independent biological replicates, each denoted by a black circle. **b)** Methanol extracts were prepared from whole mosquitoes fed on uninfected (Naïve) or *P. berghei*-infected (IBF) mice, 48 hPBF. Ecdysteroids were measured by EIA and expressed as pg/ female. Each data point represents a pool of 3-5 females. Data show the median ± interquartile range of 7-10 pools of females representing 2 mosquito generations. Between-group differences were analyzed using a Mann-Whitney test (*p*<0.0001, Mann-Whitney U=0, median naïve=146.5, median IBF=87.0, Hodges Lehmann estimated difference=-55.4, 95% CI[-43.9, -64.4]). Statistical significance is indicated as follows: *** , *p*<0.0005; **, *p*<0.01; *, *p*<0.05; ns, not significant (*p*>0.05).

### Ectopic activation of JNK signaling reduces the number of eggs developed

Since JNK proved necessary for the reproductive costs of *Plasmodium* infection we next asked whether the ectopic activation of JNK signaling might be sufficient *per se* to impair egg development. We have previously observed that JNK signaling can be induced by depletion of the JNK phosphatase *puckered (puc)* [48]. Virgin females were thus injected with a *dsRNA* targeting *puc* (*dspuc)* or *dsGFP* as a control. Injected females were allowed to take a blood meal and then, 24h PBF, levels of pJNK in the RT were measured by western blot while the number of developed eggs was counted three days PBF. Compared to *dsGFP-*injected controls, females injected with *dspuc* showed a significant decrease in levels of *puc* mRNA (**Supp. Fig. 2B**) and an increase in levels of pJNK in the RT (**Figs. 3A, 3B**) while levels of pS6K were unaffected, mirroring our observations with *Plasmodium* infection (**Figs. 1A, 1B**). This was accompanied by a decrease in egg numbers in *dspuc*-injected females relative to controls (**Fig. 3C**). In keeping with published data [48] and consistent with a dominant role for *JNK1* in this effect, both the activation of JNK signaling in the RT (**Fig. 3A**, Expt.3) and the reduction in egg number (**Fig. 3C**) were fully reversed in females in which both *JNK1* and *puc* were depleted simultaneously. As previously observed, depletion of *JNK1* alone had no significant effect on egg number. Moreover, levels of pJNK were increased in *dspuc*-injected females without impacting levels of pS6K (**Fig. 3B**). Together these data suggest that JNK signaling is sufficient *per se* to limit egg production following a blood meal.

**Fig 3.**
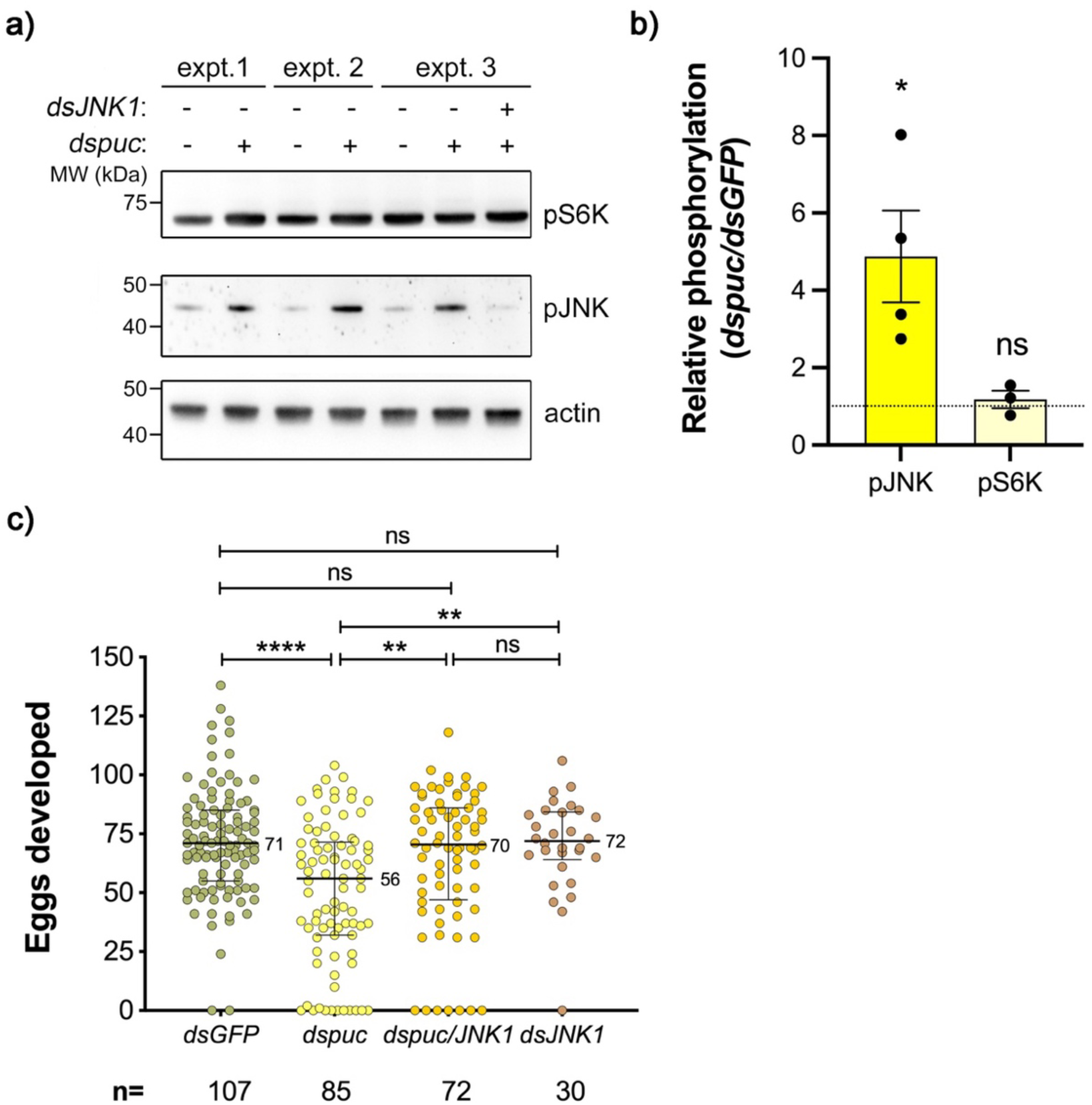
*Puckered* depletion causes JNK-dependent egg loss. **a)** A representative western blot for reproductive tracts of virgin females injected with *dspuc* (+), *dsGFP* (-) or in experiment 3, *dsJNK1* and *dspuc* and then 24 hours later allowed to take a blood meal. Reproductive tracts were dissected 24 hours PBF. Originals of the western blots presented here appear in full in Supplemantary Data**. b)** Actin-normalized densitometry values for pJNK and pS6K were used to calculate the ratio of the signal in *dsGFP*- and *dspuc*-injected females. Deviation from a ratio of 1 (indicated by the dashed line) was assessed using a using a two tailed one sample t-test (pJNK *p*=0.047, n=4, t(3)=3.26, discrepancy=3.87, 95%CI [0.096, 7.65]; pS6K *p*=0.51, n=3, t(2)=0.8, discrepancy=0.18, 95% CI [-0.79, 1.15]). The graph represents the mean ± SEM of 4 (pJNK) or 3 (pS6K) independent biological replicates, each denoted by a black circle. **c)** Virgin females were injected with *dsGFP, dspuc, dsJNK1* or *dsJNK1* and *dspuc* and then, 24 hours later, allowed to blood feed. Eggs were counted 72 hours PBF. The total number of mosquitoes analyzed (n) is reported. Data represent the median and interquartile range of 5 independent biological replicates. A Kruskal-Wallis test indicated significant differences between groups (Kruskal-Wallis statistic H(3)= 24.17, *p*<0.0001). Post hoc pairwise comparisons with Dunn’s correction revealed significant differences between *GFP* and *puc* (*Z*=4.53, p<0.0001), *puc* and *puc/JNK* (*Z*=3.42, p=0.0037) and *puc* and *JNK* (Z=3.23, p=0.0074). Statistical significance is indicated as follows: ****, *p*<0.0001; **, *p*<0.01; *, *p*<0.05; ns, not significant (p>0.05).

### JNK signaling is implicated in fitness costs induced by heat stress

We next addressed whether JNK signaling might be involved in responses to a non-infectious environmental stress such as excess heat. We first examined whether increasing the temperature at which adult mosquitoes were incubated before and after a blood meal might impact their fecundity and longevity. As shown in **Supplementary Figure 5**, virgin *An. gambiae* females incubated at 33°C for 24 hours before being blood-fed, develop significantly fewer eggs (**Supp. Fig. 5A**) and exhibit higher mortality (**Supp. Fig. 5B**) than those maintained at 27°C. These effects were not explained by alteration in the amount of blood consumed (**Supp. Fig. 5C**) and were independent of female size (**Supp. Fig. 5D**). To assess whether these fitness costs were linked to the activation of JNK signaling, we measured levels of pJNK in the RT at 24h and 48h PBF. Incubation at 33°C resulted in an increase in pJNK levels at 24h PBF compared to those incubated at 27°C and a similar but non-signficicant trend was observed at 48h PBF (**Figs. 4A, 4B**). In contrast pS6K, while strongly induced by blood feeding, was unaffected by heat stress (**Figs. 4A, 4B**). We noted the more rapid activation kinetics of both pJNK and pS6K in these experiments compared to those involving *Plasmodium* infection that we attribute to the higher incubation temperatures used here (27°C or 33°C vs 20°C, **Figs. 1A, 1B**).

**Fig 4.**
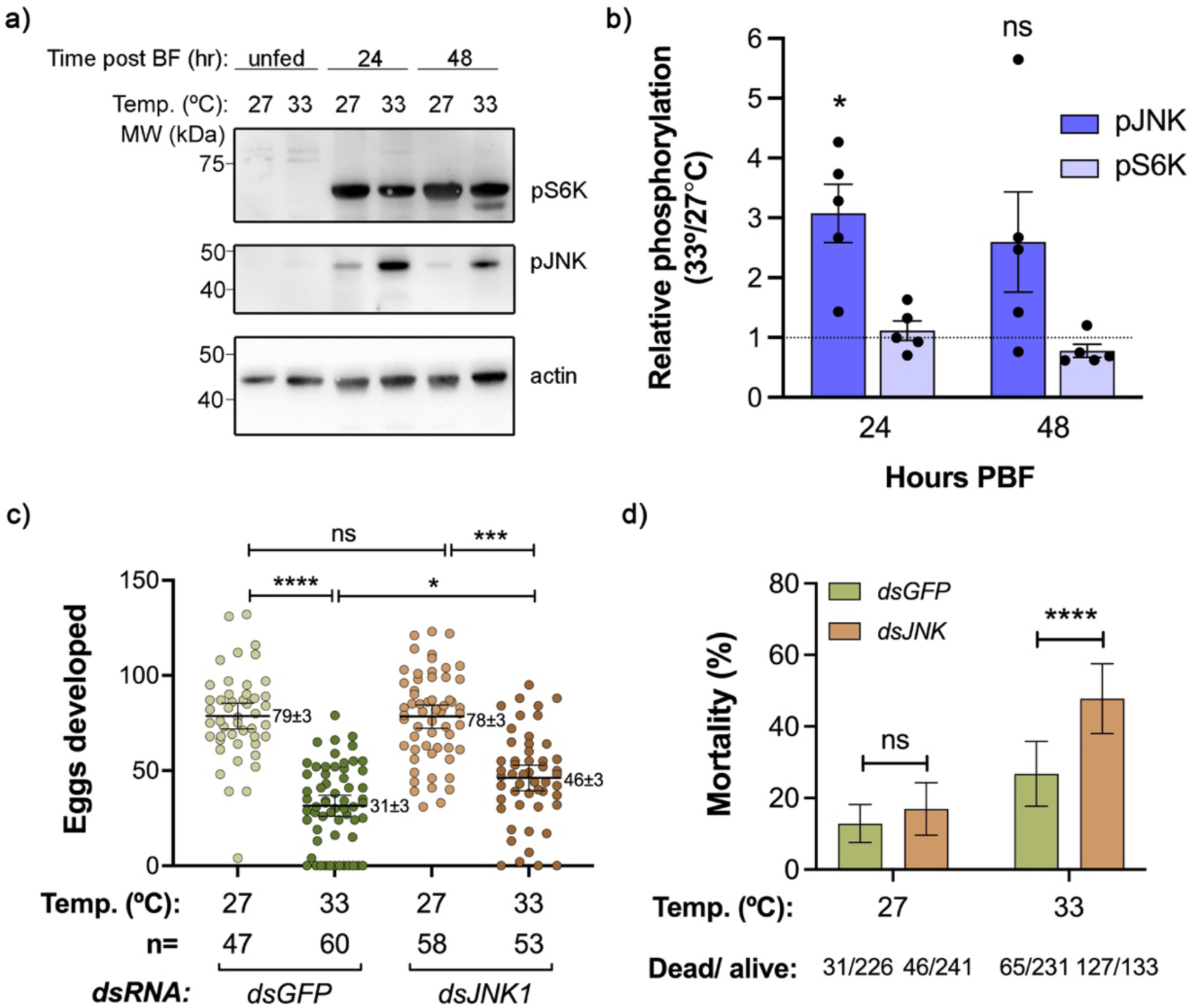
JNK contributes to heat stress-induced egg loss while promoting survival. **a)** A representative western blot for pJNK, pS6K and actin in reproductive tracts of females maintained at either 27°C or 33°C, 24 or 48 hPBF or consuming only sugar solution (Unfed). Originals of the western blots presented here appear in full in Supplemantary Data**. b)** Actin-normalized densitometry values for pJNK and pS6K were used to calculate the ratio of the signal in heat-exposed vs control females. Deviation from a ratio of 1 (indicated by the dashed line) was assessed using a using a one sample t-test. This was only significant for pJNK at 24 hPBF (p=0.013, t(4)=4.25, discrepancy=2.08, 95%CI [0.72, 3.43]). Data show the mean ± SEM of 5 independent biological replicates, each denoted by a black circle. **c)** Virgin females were injected with the *dsRNA* indicated and 24 hours later placed at 27 °C or 33°C then, after a further 24 hours, allowed to blood feed. Eggs were counted 72 hours PBF. The total number of mosquitoes analyzed (n) in each group is indicated. Data represent the mean ± 95%CI of 3 independent biological replicates. The mean ± SEM is also shown numerically. Differences in egg number were compared using a negative binomial generalized linear model with Tukey’s correction of pairwise comparisons (see Methods). Selected comparisons are indicated: *dsGFP* 27/*dsGFP* 33 (effect ratio=2.5, 95%CI [1.78, 2.37], Z= 6.96, *p*<0.0001); *dsJNK1* 27/*dsJNK1* 33 (effect ratio=1.7, 95%CI [1.22, 2.37], Z=4.1, *p*=0.00024; d*sGFP* 33/*dsJNK1* 33 (effect ratio=0.68, 95%CI [0.49, 0.95], Z=-2.96, *p*=0.016). (**d)** *dsGFP-* or *dsJNK1*-injected females were placed at either 27°C or 33°C and 24 hours later allowed to take a blood meal. Mortality over the first two days PBF was monitored and calculated as a percentage of the total in each of 6 independent biological replicates. The mean % mortality ± SEM is presented in the graph and the total number of dead vs live mosquitoes is reported below. Differences in the mortality rate between *dsGFP-* and *dsJNK1*-injected females at the two different temperatures were compared using Fishers exact test on the absolute values. The odds ratio of survival between *dsGFP-* and *dsJNK1* was significant at 33°C (*p*<0.0001, OR=3.39, 95%CI [2.44, 4.57]) but not at 27 °C (*p*=0.22, OR=1.39, 95%CI [0.85, 2.28]). Statistical significance is indicated as follows: **** *p*<0.0001; **, *p*<0.01, *, *p*<0.05, ns, not significant (*p*>0.05).

To establish whether *JNK1* was required for the reduced fecundity associated with heat stress, females were injected with *dsGFP* (controls) or *dsJNK1* then exposed or not to heat stress (33°C). *dsJNK1* had no effect on egg number in unstressed females, as previously observed (**Figs. 1C, 3C**). In contrast, *JNK1* depletion significantly reduced levels of pJNK in the RT (**Supp. Figs. 6A, 6B**) compared to controls, and partially rescued egg production (**Fig. 4C**), but significantly increased mortality (**Fig. 4D**), under conditions of heat stress. Collectively, these data show that despite little impact on growth-linked signals, heat stress activates JNK signaling that reduces egg numbers while promoting survival.

### Heat stress or dspuc reduce 20E levels after blood feeding

Given their shared capacity to activate JNK signaling in the RT we then tested whether, like *Plasmodium* infection, *dspuc* treatment and heat stress might limit egg production by reducing the amount of 20E produced after a blood meal. We did this in two ways. First, to examine whether the activation of JNK was sufficient *per se* to suppress 20E production in the context of a blood meal, we analysed ecdysteroid levels in blood-fed control females (injected with *dsGFP*) or *puc*-depleted females, this time using MS/MS to quantify all ecdysteroids individually [56]. The levels of ecdysone, its derivative 20E and their respective 22-phosphorylated forms were all significantly reduced in the *puc*-depleted females. Secondly, having shown that *puc* depletion impacts multiple ecdysteroids, we exposed females to high temperature (33°C) and used an EIA kit that measures a range of ecdysteroids, to assess the impact of heat stress on ecdysteroid production after a blood meal. Akin to the effects of *P. berghei* infection (**Fig. 2B**), *dspuc* (**Fig. 5A**) and heat stress (**Fig. 5B**) led to significantly reduced levels of ecdysteroids compared to controls, again at a time point coincident with the peak in JNK activation PBF. Together, these data support the hypothesis that the biochemical (*dspuc*) or environmental (*P. berghei* infection, heat stress) activation of JNK signaling attenuates egg development through a shared mechanism; inhibition of blood feeding-induced ecdysteroid production and thus hint at a generalisable regulatory circuit through which environmental stress constrains reproductive investment in *An. gambiae*.

**Fig 5.**
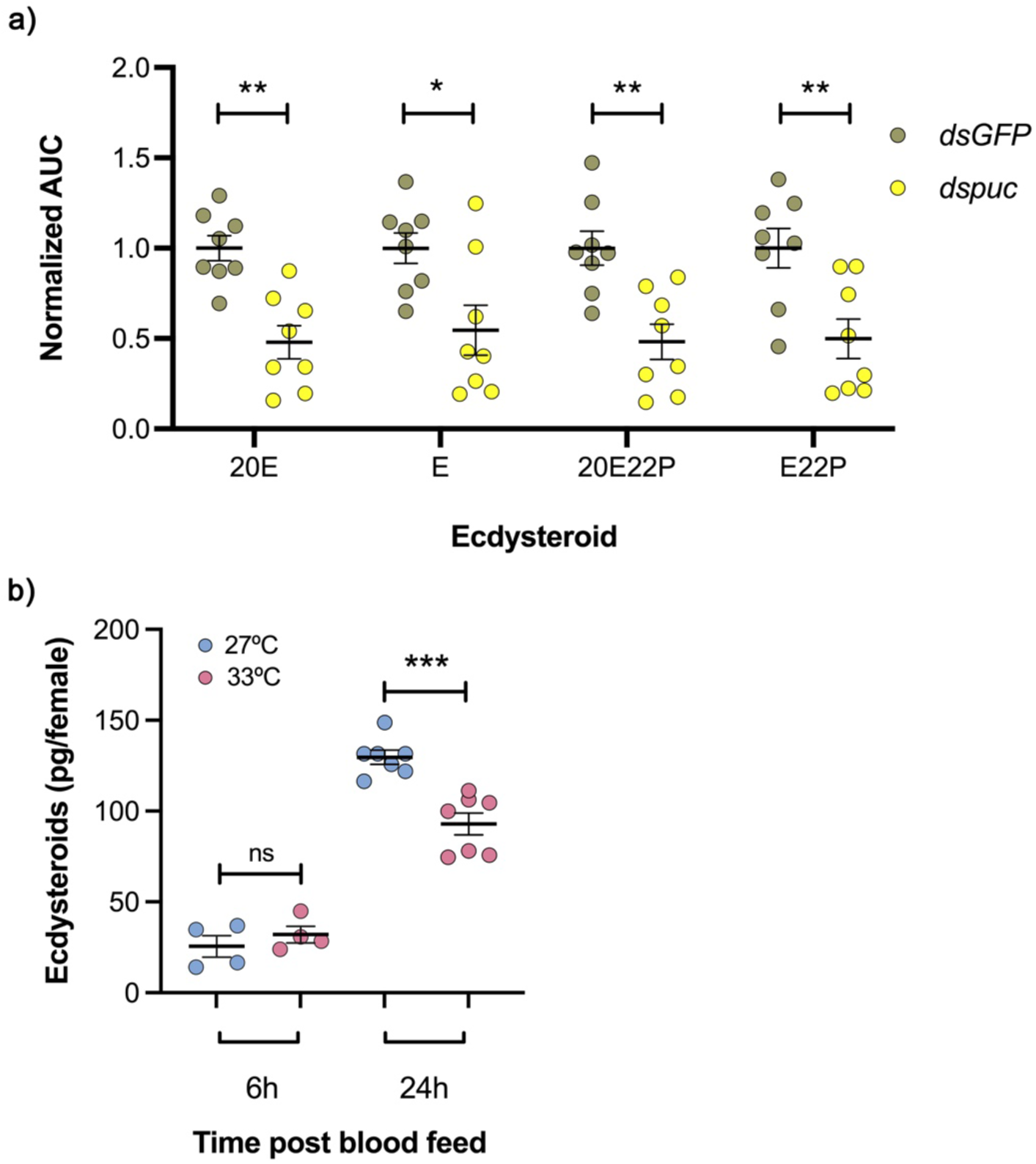
*Puckered* depletion or heat stress inhibits ecdysteroid production after blood feeding. **a)** *dsGFP-* or *dspuc*-injected females were blood-fed and the following ecdysteroids recovered in methanol extracts of whole females 24h PBF, were measured using MS/MS: 20-hydroxyecdysone (20E), ecdysone (E), and the respective 22-phosphorylated forms (20E22P and E22P). For each ecdysteroid, the mean normalized area under the curve (AUC) value in *dspuc*-treated females was expressed relative to the mean value in *dsGFP*-injected control females which was given an arbitrary value of 1. Data show the mean ± SEM of 8 pools of 50 females derived from 5 biological replicates. Between-treatment differences were analyzed using individual unpaired t tests on the unnormalised data followed by false discovery rate correction (20E, t(14)=4.54, pFDR=0.0023, *dsGFP* mean=1.84, mean difference=-0.96, 95%CI [-1.41, -0.51]; E, t(14)=2.81, pFDR=0.0172, *dsGFP* mean=1.78, mean difference=-0.81, 95% CI [-1.43, -0.19]; 20E22P, t(14)=3.85, pFDR=0.0037, *dsGFP* mean=0.30, mean difference=-0.16, 95% CI [-0.24, -0.069]; E22P, t(14)=3.26, pFDR=0.0080, *dsGFP* mean=0.49, mean difference=-0.25, 95% CI [-0.41, - 0.084]) **b)** Methanol extracts were prepared of whole mosquitoes exposed (33°C) or not (27°C) to heat stress at the indicated time points PBF. Ecdysteroids were measured by EIA and expressed as pg/female. Each data point represents a pool of 3–5 females. Data show the mean ± SEM range of 4–7 pools of females representing 3 mosquito generations. Between-group differences were analyzed using an unpaired t test (6h, mean difference=6.47, 95%CI [-11.83, 24.77], t(6)=0.87, p=0.42); 24h, mean difference=-36.74, 95%CI [-52.39, -21.09], t(12)=5.11, p=0.0003). Statistical significance is indicated as follows: *** , *p*<0.0005; **, *p*<0.01; *, *p*<0.05; ns, not significant (*p*>0.05).

## DISCUSSION

Here, we use the malaria vector *Anopheles gambiae* to understand how nutritional resources are divided between the competing energy demands of reproduction and survival. The costs of infectious and environmental stresses on mosquito fitness, and specifically reproductive output, are well documented [17–19,25] and we explain how these effects are realized through JNK signaling, revealing that some details appear conserved between distinct stressors.

The model supported by our findings, summarised in **Figure 6**, suggests that even in the absence of deliberately applied stress, blood feeding is linked to an increase in JNK activity in the reproductive tract, possibly downstream of documented increases in reactive oxygen species [57] and bacterial proliferation [58] which promote JNK signaling [59,60].

**Fig 6.**
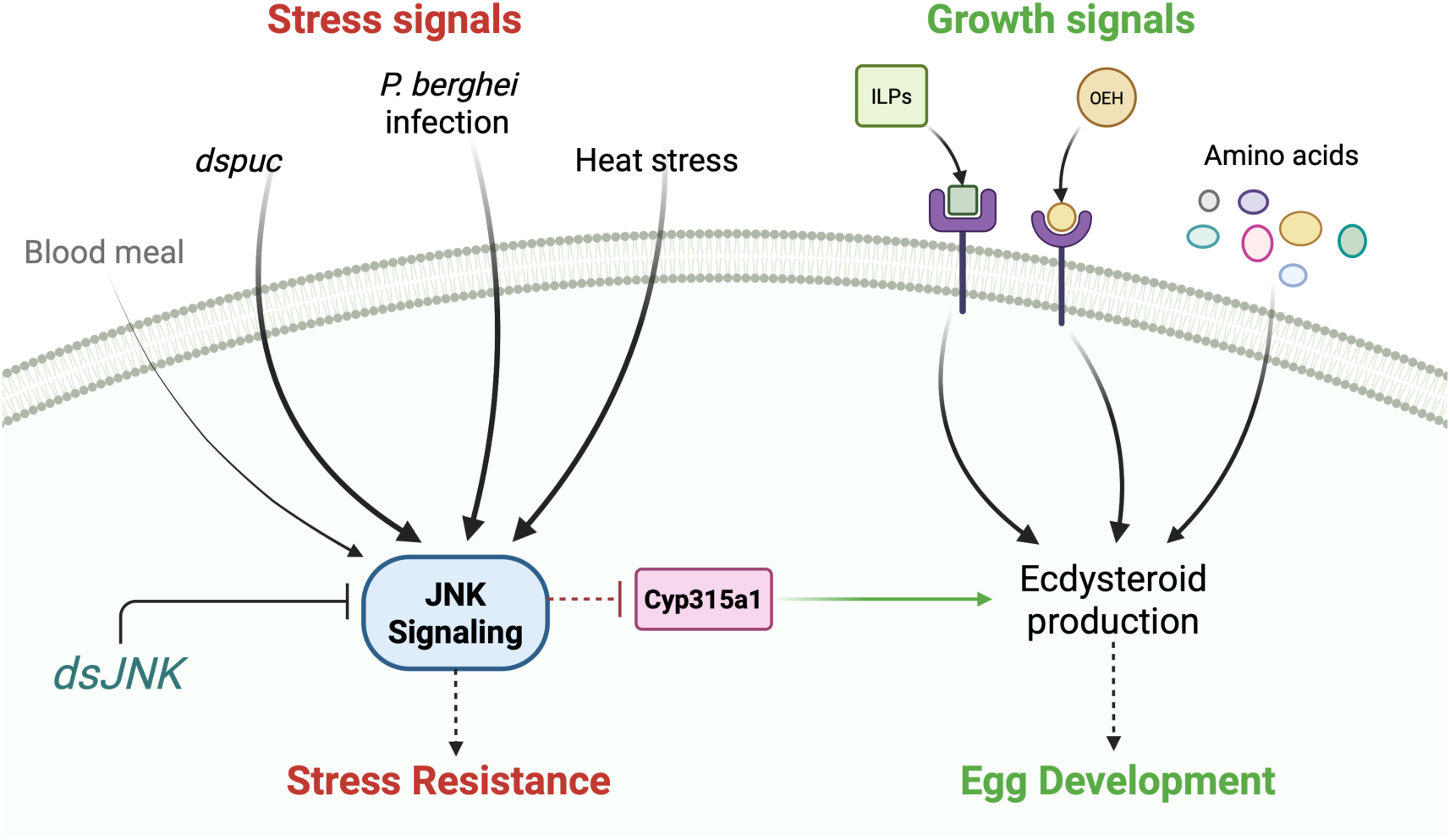
Schematic representation of findings. After a blood meal in the absence of stress signals, JNK signaling is weak (indicated by the fine arrow) and growth signals comprising Insulin-like peptides (ILPs), ovarian ecdysteroid hormone (OEH) and free amino acids, predominate. This drives the production of ecdysteroids and robust egg development. After a blood meal in the presence of physiological (*Plasmodium* infection), biochemical (*puc* depletion,) or environmental (heat) stress stimuli, JNK is over-activated in the reproductive tract promoting stress resistance but also suppressing ecdysteroid production and consequently egg production. In JNK-depleted females, JNK activity is reduced, favouring expression of *Cyp315a1* resulting in increased egg production under stress. Figure created in https://BioRender.com.

The boost to this JNK signal when the blood meal takes place in the context of elevated stress (*P. berghei*-infection [**Fig. 1C**], *puc* depletion [**Fig. 3C**] or high temperature [**Fig. 4C**]), is necessary and sufficient to attenuate mosquito fecundity. In a possible echo of the link between stress-induced reproductive costs and sex steroid production in mammals [1], we connect this effect mechanistically to the JNK-dependent suppression of ecdysteroid production that drives oogenesis in this species.

The key elements of this model are well supported in the literature. The activation of JNK by stressors including infection and elevated temperature [30,31] and the negative impact of those stressors on egg development [17,25] have been reported. In the specific case of *Plasmodium* infection, fitness costs of infection are correlated with JNK activation; *P. berghei* infection engages the JNK pathway [24] and causes significant fitness costs to the mosquito host [19]. In contrast, *P. falciparum*, likely thanks to its JNK-silencing *Pfs47* alleles [22], fails to engage the JNK pathway [23] and imposes no such fitness costs [20]. Our data build on this correlation to establish a causal link between JNK activation and *P. berghei*-associated fitness costs. Notably, previous reports have documented a role for JNK in limiting *P. berghei* infection with *JNK* depletion causing an increase in oocyst numbers [24]. We observed a similar effect but only in three of six experiments where control infection intensity was consistent with those studies (<10 oocysts/midgut); at higher infection intensities that link was lost (**Fig. 1, Supp. Fig. 3**). This is in line with other reports of infection intensity-dependent effects of depletion of immune-related pathways [61,62] as well as qualitative differences in transcriptional responses to high and low intensity *Plasmodium* infections [63,64] and could be due to insufficient statistical power in more variable high-intensity infections.

A second discrepancy between our data and previous studies [22,24] was the observation of a decrease in infection prevalence in JNK-depleted females, i.e. a larger number of *dsJNK1*-treated females who fed on infected blood, killed all the parasites within it. Notably, the same effect was not seen in the low intensity infections suggesting that the reduced prevalence is associated specifically with the higher intensity infections that distinguish our work from others [22,24]. Moreover, if the fitness costs of infection are caused by immunopathology (as discussed below), it may be the females who responded most robustly to the parasite (by clearing the infection) whose survival is favoured by the relative absence of JNK.

The targeting of ecdysteroid synthesis as a means to reallocate resources between reproduction and immunity in insects has been proposed previously [65] and, as alluded to above, is reminiscent of stress-induced changes in sex steroid production in mammals [1]. The importance of *Cyp315a1* in particular is consistent with its upregulation in blood-fed mosquitoes [66] and selective downregulation in bacterially-infected or wounded mosquitoes [67], insults that activate JNK signaling [28]. Similarly, in other insects in which ecdysteroids regulate reproduction, stressful (diapause-inducing) environmental cues suppress expression of *Cyp315a1* [68,69] while its experimental depletion or mutation phenocopies our results, reducing 20E production and reproductive output [70,71].

How might JNK signaling inhibit ecdysteroid synthesis? Insulin-like peptides are important drivers of ecdysone synthesis after a blood meal [9] while JNK antagonises insulin signaling [35,37,39,40]. Thus, our observations could be explained by stress-induced activation of JNK reducing insulin production in the brain, or its signaling in the periphery, both of which have been documented [35]. However, reduced insulin signaling should be reflected in reduced levels of pS6K which we do not observe. An alternative is that JNK could activate the transcription factor FoxO [34,37] which inhibits insulin signaling at the transcriptional level, without impacting levels of pS6K [35,37]. Interestingly, FoxO has a pivotal role in diapause in several genera including mosquitoes [72]. It will be interesting in future experiments to examine the involvement of FoxO in the stress-induced reduction in eggs we observe, as well as a possible role for JNK in diapausing mosquitoes.

Implicit in the idea of a trade-off between reproduction and survival is that over commitment to one should come at the cost of the other, as others have described [73,74]. In the case of heat-stress our data are consistent with such a prediction; JNK depletion increases reproductive investment at the cost of increased mortality, presumably because the JNK pathway increases tolerance to heat stress, for example through the production of heat shock proteins [31,32].

In contrast, for *P. berghei* infection, the lack of JNK simultaneously rescues the number of eggs developed and reduces mortality. It is possible that JNK deficiency in the context of *P. berghei* infection may impose costs that we are not measuring. However, the reduced mortality in the case of JNK depletion supports a growing body of evidence that the fitness costs of *P. berghei* infection do not result from parasite infection *per se.* Depending on the model used, *Plasmodium* infection may impose limited costs [16,19,20] or even benefit the mosquito [75]. Rather, costs to the host appear to stem from ‘immunopathology’ caused by the immune response to the parasite, akin to those resulting from the unconstrained activation of the Toll pathway [76]. Our findings thus make clear that the evolution of JNK-silencing *Pfs47* alleles described in *P. falciparum* [22,23] would have benefited both mosquito and parasite: the parasite promotes its own survival in low level infections whilst reducing the chances of its vector’s premature death, favouring transmission. At the same time, by reducing the longevity and fecundity costs of infection, the lifetime reproductive output of the infected mosquito is protected.

Taken together, the data presented here support a causal link between stress-induced activation of JNK in the RT and a reduction in fecundity via the suppression of ecdysteroid synthesis. In the case of heat stress, JNK appears to mediate a trade-off between egg development and survival. In contrast, the JNK-dependent immune response to the non-adapted parasite *P. berghei* is revealed to be significantly effective only at low infection intensities whilst also being detrimental to mosquito fitness, reducing egg numbers and increasing mortality. Indeed, by limiting the JNK response, *P. falciparum* may boost its own survival as well as its transmission by promoting the longevity and reproductive fitness of its natural mosquito host, highlighting the critical role of coevolution between vector and parasite in malaria transmission.

## METHODS

### Ethics statement

Experiments with animals were performed in line with the EU Animals Act 1986. *Plasmodium berghei* (*P. berghei*) infection of mosquitoes by blood feeding on infected mice was approved by the Italian Ministry of Health (License # 1184/2020-PR) based on the D. lgs. 26/2014. Procedures used were designated mild-to-moderate severity. All efforts were made to minimize the numbers of animals used, while protocols were kept under review to reduce, refine, and replace animal usage wherever possible.

### Mosquito rearing

Mosquitoes were reared in a level 2 containment facility in the Department of Medicine and Surgery, Perugia, Italy, authorization N. PG/IC/Imp2/13/001-Rev2 from the Italian Ministry of Health. *An. gambiae,* G3 strain (MRA-112, MR4/BEI resources) were maintained at ∼28°C and 60–80% humidity with a 12/12 h day/night cycle. Larvae were cultivated according to the MR4 protocol (https://www.beiresources.org/Portals/2/VectorResources/2016%20Methods%20in%20Anopheles%20Research%20full%20manual.pdf). Adults were fed *ad libitum* on 6% glucose supplemented with methylparaben (0.1%) (Sigma-Aldrich). Females were blood fed weekly with bovine blood using a Hemotek PS5 membrane feeder system (Discovery Workshops, UK). Experimental females were separated from males at the pupal stage by microscopic examination of the terminalia and retained as virgins. In experiments to examine the effect of *P. berghei* infection on fecundity and fertility of mated females, females and males were maintained in the same cage (mixed cage) for 3 nights before feeding on a *P. berghei*-infected or uninfected mouse. Females were transferred to individual oviposition cups after blood feeding and the number of eggs laid checked each day. In females failing to oviposit at d8 PBF, mating was verified by microscopic detection of sperm in the spermatheca. Wing length was measured on occasion by microscopic examination of slide-mounted pairs of dissected wings using Image J 1.54g software. Wing length was taken to be the mean distance from the aula notch to the wing tip for each pair [77]. The average blood meal size was calculated by measuring the mean mass of randomly selected groups of 10–30 females before and after blood feeding.

### P. berghei infections

*P. berghei* infections were performed essentially as previously described [78]. *P. berghei* parasites (5×10^6^) expressing GFP (Bergreen, [79]) were injected into CD1 mice (Charles River, Sant’Angelo Lodigiano, Italy) by the intraperitoneal route. Three days later, parasitemia was monitored by Giemsa staining (Hemacolor Rapid staining: Merck/ Millipore, Milan, Italy) of tail blood smears to verify gametocyte levels of 0.8–1.5%. Mice were anesthetized by intraperitoneal injection with ketamine and xylazine (100 mg/kg and 40 mg/kg, respectively). *An. gambiae* females were then allowed to feed (30 min) on a *P. berghei*-infected mouse or an uninfected littermate control and placed at 20°C to allow *P. berghei* development. Females that failed to blood feed were removed from the cage. In RNAi experiments, mosquitoes were injected 2 days before blood feeding with *dsRNA*s targeting *JNK1* (*dsJNK1*) or *GFP* (*dsGFP*) as a control. Infection intensity and egg development in each female were assessed eight days post infection (d8 PI) by counting the number of eggs in reproductive tracts and fluorescent oocysts in midguts (Evos FL Imaging System AMG). Females fed on an infected mouse but failing to develop oocysts were excluded from the egg number and infection intensity counts but included in the calculation of infection prevalence. In experiments (RNA-seq, western blotting or ecdysteroid measurement) where the time point analyzed preceded verifiable infection, some mosquitoes from the same cohorts were retained for subsequent analysis at d8 PI. In some experiments (Supp. Fig. 1D) mosquitoes were fed on an infected or uninfected mouse as above then maintained at either 20°C (allowing *P. berghei* development) or 27°C (precluding *P. berghei* development). The number of eggs developed and infection intensity in all three conditions was measured at d8 PI.

### RNAi experiments

The design, production and injection of the *dsRNA*s used in this study have all been described in detail previously [48]. Briefly, 3–4-day-old virgin females were injected (Nanoject II, Drummond Scientific/Olinto Martelli Srl, Florence, Italy) with 0.69μg (138nl, 5μg/μl *dsRNA*s) targeting *JNK1* (*dsJNK1*), *puckered* (*dspuc*) or on occasion, both *dsJNK1* and *dspuc* (prepared such that 138nl contains 0.69μg of each *dsRNA*). A *dsRNA* targeting *GFP (dsGFP)*, a gene not expressed in these mosquitoes, was used as a negative control [47]. Assignment of mosquitoes to different groups was random but not blinded. To examine the effects of heat or *P. berghei* infection stress on fecundity, mosquitoes were allowed 48 hours post injection (HPI) to recover before being blood fed. In *dspuc* experiments, females were blood fed 24 HPI. The number of fully developed eggs per female was counted 72 hours post blood feeding (hPBF). The knock down efficiency of each *dsRNA* was defined by measuring the expression of the gene of interest by qRT-PCR (see below) in females in which the gene was targeted relative to that in *dsGFP*-injected controls using whole unfed females at 48 HPI (*dsJNK1*) or 24 HPI (*dspuc*).

### Heat stress experiments

Three to four-day-old virgin females from the same generation were placed in an incubator at 33°C (heat stressed) or 27°C (unstressed) for 24 hours before and 72 hPBF. Females that failed to blood feed were removed from the cage. Daily survival rates were assessed and the number of eggs developed counted 72 hPBF. In experiments to examine the impact of JNK1 on responses to heat stress, females were injected with *dsJNK1* or *dsGFP* as described above then allowed 24 hours to recover before being placed at the appropriate temperature for a further 24 hours before blood feeding.

### Western blotting

Reproductive tracts (comprising ovaries, atrium and spermatheca) were recovered from 10-12 virgin females maintained only on sugar (unfed) or at various time points after blood feeding and prepared and analyzed as previously described in detail [48] with minor modifications. Tissues were dissected in to 30µl of extraction buffer containing 1% SDS, rather than 0.1% described previously [48], then 10µl of 4x SDS-sample buffer (Thermofisher) supplemented with 20mM DTT was added prior to manual homogenization with a pestle, clarification by centrifugation (14000 x g, 5 min, RT) and denaturation at 85°C for 5 min. Tissue homogenates were resolved using precast 4-12% bis/tris gels (Thermofisher) and transferred and blocked as described elsewhere [48]. Blocked membranes were cut at the 60 kDa marker and blotted (O/N, 4°C, 1/1000 in 5% BSA) with either anti-pJNK (MW <60kDa, Cell Signaling Inc, RRID: AB_823588) or rabbit anti-pS6K (MW >60kDa, Merck/Millipore, Cat. No. 07-018-L). The lower half of the membrane was stripped (15 min, RT, ReStore; Thermofisher) and reprobed (1h, 1/5000 in TBS-tween with 4% milk powder) with anti-actin (AbCam, Cambridge, UK, RRID: AB_867488). HRP-conjugated anti-rabbit (RRID: AB_2536530, pJNK, pS6K) and anti-rat (RRID: AB_228356, actin) secondary antibodies (Thermofisher) were detected by the addition of enhanced chemiluminescence (ECL) reagents (Amersham, Cambridge, UK) and visualized using an iBright CL1500 imaging system (Thermofisher). Optical density of bands was measured using ImageJ software [80], normalizing above-background values for pJNK and pS6K to the actin values in each lane. This ratio (pJNK/actin and pS6K/actin) was used as a measure of ‘relative phosphorylation’ for each kinase in each sample. Between-sample differences in relative phosphorylation were compared using a one sample t test, testing the null hypothesis that the relative pJNK signal in ‘stressed’ females (*P. berghei*-infected, heat-stressed, *dspuc-*treated) vs ‘unstressed’ females (control) should be the same.

### Gene expression analysis by qRT-PCR

Gene expression was assessed by qRT-PCR as described previously [48,81]. Briefly, dissected tissues were recovered directly to 10μl of RNA-later (Ambion), immediately supplemented with 250μl Tri-reagent and homogenized using a motorized pestle. RNA in supernatants clarified by centrifugation (14000 x *g*, 15 min, 4°C) was extracted, DNase digested and eluted using Direct-zol RNA miniprep columns (Zymo Research/ Euroclone, Milan, Italy) according to the manufacturer’s instructions. Some of this material (0.5–1μg) was reverse transcribed to cDNA as described previously [82]. Triplicate 5μl aliquots were analyzed using Fast SybrGreen master Mix (Thermofisher) and the forward and reverse primers listed in **Supplementary Table 3**. Reactions were run on a QuantStudio 3 thermocycler (Thermofisher) and differences in expression between treatments were quantified using the delta delta CT method. Expression for each gene of interest was first calculated relative to that of a blood feeding-insensitive reference gene (*Ribosomal protein L19*, *RpL19*; AGAP004422) in the same sample [83].

### RNA-seq

Three to four-day-old females were injected with *dsJNK1* or a control (*dsGFP)* and after 48 hours, allowed to feed on a *P. berghei*-infected mouse (see above) or a naïve littermate control. Twenty-eight hours later, reproductive tracts (ovaries, atrium and spermatheca) from 15-20 females were recovered and stored in Trizol (250µL) supplemented with RNAlater (25µl). RNA was recovered from tissue homogenates using Direct-zol RNA miniprep columns (Zymo Research/ Euroclone, Milan, Italy) as described above, eluted in 18µl of H_2_O and stored at -80°C. Each experiment comprised 4 conditions: *dsGFP,* naïve blood feed; *dsGFP,* infected blood feed (IBF); *dsJNK1,* naïve; *dsJNK1,* IBF. Five biological replicates comprising 20 samples were analyzed for RNA quality by PoloGGB, Siena, Italy (https://www.pologgb.com) using a fragment analyzer (Agilent) on the basis of which, one biological replicate (Experiment 1) was abandoned. Illumina Nextseq 550 paired-end libraries were prepared from the remaining 16 samples using a QIAseq stranded mRNA kit (QIAGEN, Milan, Italy) and run by PoloGGB. The resulting FastQ files are available through the European Nucleotide Archive: https://www.ebi.ac.uk/ena/browser/view/PRJEB35279. Bioinformatic analysis and preparation of volcano plots was performed using the online Galaxy portal (https://usegalaxy.org) according to the accompanying online tutorials. Quality control of sequencing reads was performed using FastQC (Galaxy version 0.74+galaxy1) and trimmed using Cutadapt (Galaxy version 4.9+galaxy1). Trimmed sequencing reads were aligned to the *An. gambiae* genome assembly idAnoGambNW_F1_1 (https://www.ncbi.nlm.nih.gov/datasets/genome/GCF_943734735.2/) and the number of reads mapped to genes was counted using using RNAstar (Galaxy version 2.7.11a+galaxy1). Normalized read counts were calculated and differential gene expression analyzed using DESeq2 (Galaxy version 2.11.40.8+galaxy0). Genes listed as differentially expressed (**Supp. Tables 1 and 2**) were those passing a multiple comparison-adjusted statistical threshold of p<0.05. No minimum ‘fold-change’ threshold was applied.

### Ecdysteroid measurements. MS/MS

Ecdysteroids in methanol extracts of whole females were measured using tandem mass spectrometry as previously described [56]. Whole mosquitoes (50 per point) were homogenized in 100% methanol (2ml) using a bead beater (2-mm glass beads, 2,400 r.p.m., 90 s). A second methanol extraction was performed on the pellet and pooled methanol extracts were clarified by centrifugation, dried under nitrogen flow and resuspended in 80% methanol in water (50 µl). Samples were resolved by liquid chromatography (LC, Vanquish, Thermo Fisher) before being analyzed on an ID-X mass spectrometer (Thermo Fisher) as described in detail elsewhere [56].

### EIA (Enzyme Immunoassay)

In some experiments ecdysteroid levels were measured using an EIA kit (Cayman Chemical, Vinci Biochem, Florence, Italy) according to the manufacturer’s instructions and as previously described [16,82]. Briefly, pools of 3–5 whole females were homogenized in methanol using a motorized pestle. Methanol extracts were allowed to evaporate to dryness and resuspended in reaction buffer. Absorbance at 405nm upon addition of the substrate (Ellman’s reagent), inversely related to the quantity of ecdysteroid in the sample, was measured using a plate reader (BMG Labtech spectrostar nano, Euroclone Spl. Milan, Italy) and quantified by reference to a standard curve using Prism v10.3.1 software.

### Statistics and Reproducibility

Statistical analyses were performed using Prism 10.3.1 (GraphPad, La Jolla, USA) or on occasion Rstudio v2025.05.0+496 (Posit Software, Boston, USA). Non-parametric methods were used unless normality of the data could be demonstrated using a Shapiro-Wilk test. Where parametric tests were used the mean ± standard error of the mean (SEM) are presented while for non-parametric tests the median ± interquartile range (IQR) or 95% confidence intervals (95% CI) are presented. In all tests, statistical significance was ascribed to *p*<0.05. Differences between two groups were assessed using Student’s t test (parametric) or Mann-Whitney (non-parametric) tests. In all cases, two-tailed tests were performed. Differences between multiple treatment groups were assessed using one-way ANOVA tests with Dunnett’s correction (for parametric tests) or Kruskal-Wallis tests with Dunn’s correction (for non-parametric tests). Where two categorical variables were present but egg counts were not normally distributed (**Figs. 1c and 4c**), data were analysed using a negative binomial generalized linear model (GLM) in Rstudio to account for overdispersion and the presence of zero counts. Model fit was assessed using Pearson residuals and a chi-square goodness-of-fit test, confirming that the negative binomial GLM appropriately accounted for the variance in the dataset. Treatment (*dsGFP* vs *dsJNK1*), stress (temperature or *P.berghei*-infection), and their interaction were included as fixed factors. Estimated marginal means and 95% confidence intervals were calculated from the model, and pairwise comparisons between treatment × stress groups were performed using the emmeans package in R.

Between group differences in death rates were analyzed using a Mantel-Cox Log-rank test. The correlation between egg number and wing length was investigated using a Pearson’s linear regression analysis and the difference in y intercept between unstressed and heat-stressed females was compared using a t test. In western blotting and some qRT-PCR experiments the effect of a treatment (e.g. *dsJNK1* vs *dsGFP*) was assessed using a one sample t-test where normality of all groups passed a Shapiro-Wilk test or a Wilcoxon signed rank test where one or more groups failed (**Fig. 2a**). The number of replicates of each experiment performed is indicated in the legend to each Figure. Unless otherwise stated, ‘replicate’ refers to the same experiment performed on mosquitoes taken from different generations of the same G3 mosquito line.

## Supporting information

Contains Supplementary Figures 1-6, Supp. Tables 1-3 and original western blots

