## Supplementary material for "JNK signaling regulates reproductive trade-offs after *Plasmodium* infection in the malaria mosquito": Contains Supplementary Figures 1-6, Supp. Tables 1-3 and original western blots

### Current adress: Department of Microbiology and Immunology
University of Texas Medical Branch, Galveston, TX, USA

† Current adress: University of Ostrava, Ostrava, Czechia

§ Current adress: Center for Global Infectious Disease Research, Seattle Children's Research Institute, Seattle, WA, USA

¶ Current adress: Regel Therapeutics, Cambridge, MA, USA

* Corresponding Authors
 (MJP)

 (RS)


**This PDF file includes:**

Supplementary Figures S1 to S6
Supplementary Tables S1 to S3 and legends
Supplementary Data: Original annotated western blots
Supplementary References

#### Supplementary Information

##### Supplementary Figure 1

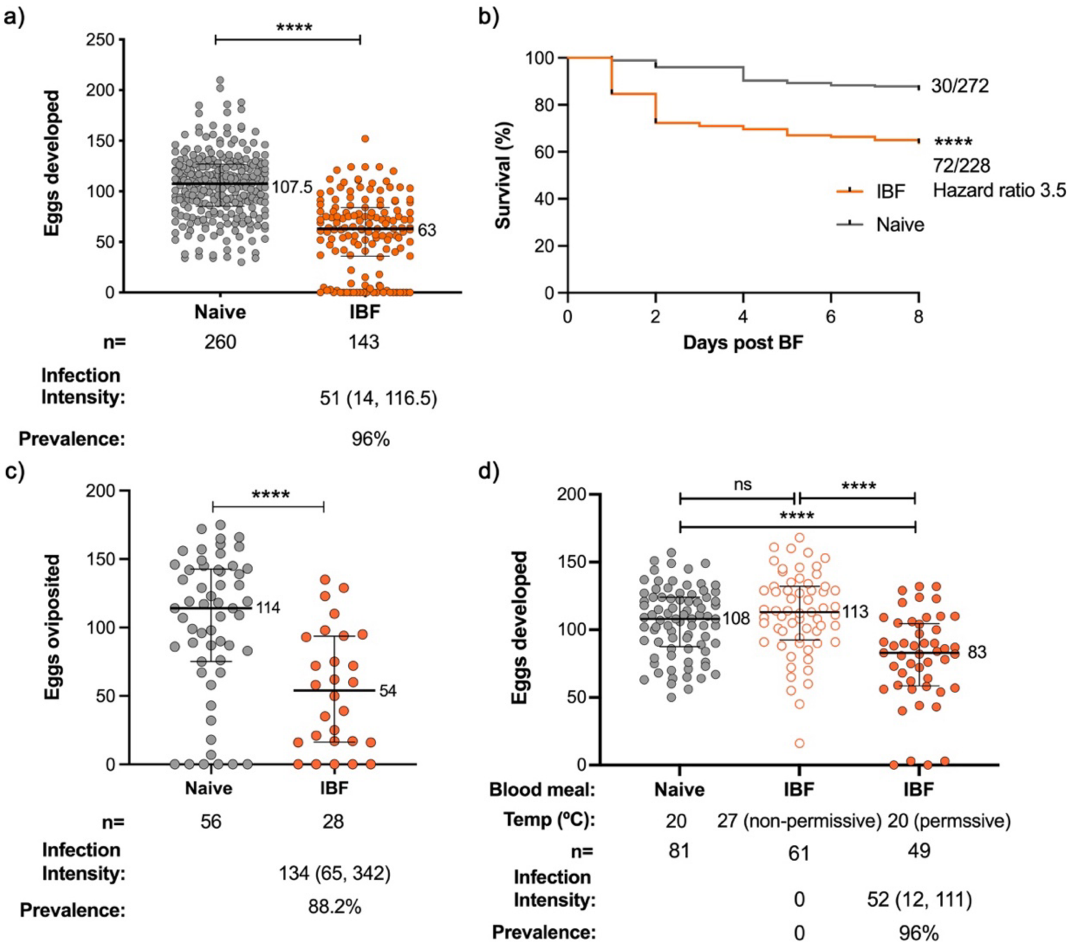


**Supplementary Figure 1. *P. berghei* infection decreases mosquito fecundity, survival and oviposition. a).** Virgin females were fed on an uninfected (Naïve) or *P. berghei*-infected mouse (IBF) and eggs and parasites counted at day 8 PBF. Data show the median ± interquartile range of 8 biological replicates. Differences in fecundity were analyzed using a Mann-Whitney test (Mann-Whitney U=6328, p<0.0001, median Naïve=107.5, median IBF=63, Hodges-Lehmann estimate=45, 95% CI [38.0, 53.0]). The number of mosquitoes analyzed (n), infection intensity (median oocysts/midgut with interquartile range) and prevalence (% of blood-fed females containing parasites) are shown. **b)** Data show daily survival rates of virgin females up to day 8 following an uninfected (Naïve) or *P. berghei*-infected blood meal (IBF) from 4 independent biological replicates and presented as % surviving females on each day. The numbers of dead/total females are also reported. Differences in survival were analyzed using a Log-rank (Mantel-Cox) test and the relative hazard ratio (HR, the relative risk of death in IBF vs Naïve groups) calculated; Chi square=40.24, df=1, p<0.0001, HR=3.49, 95%CI [2.35, 5.18]. **c)** Mated females were fed on a *P. berghei*-infected (IBF) or an uninfected (Naïve) mouse then allowed up to 8 days to oviposit. Data show the median ± interquartile range of eggs oviposited in two independent biological replicates. The number of oviposited eggs was compared with a Mann-Whitney test (Mann-Whiteny U=377, p<0.0001, median naïve=114, median IBF=54, Hodges-Lehmann estimate=--53.5, 95% CI [-27.0, -57.0). The number of mosquitoes analyzed (n) is indicated. Infection intensity, and prevalence in those females surviving to d8 post BF (n=15) is reported. **d)** Virgin females were fed on a *P. berghei*-infected (IBF) or an uninfected (Naïve) mouse, then maintained at 20ºC or 27ºC, respectively permiting or precluding infection. Eggs and oocysts were counted at d8 PBF. The number of mosquitoes analyzed (n) is indicated. Infection intensity and prevalence is reported for all females at 27ºC and a representative sample (n=26) of females at 20ºC. Differences in egg number were compared using a Kruskal-Wallis test ($H(2) = 29.62$, $p < 0.0001$). Dunn's post-hoc pairwise comparison revealed significant differences between IBF20 and naïve ($p<0.0001$) and IBF20 and IBF27 ($p<0.0001$), whereas naive and IBF27 showed no significant difference ($p=0.61$). The data shown represent the median ± interquartile range of three independent biological replicates. In all tests, **** denotes $p<0.0001$; ns, not significant ($p>0.05$).

#### Supplementary Figure 2

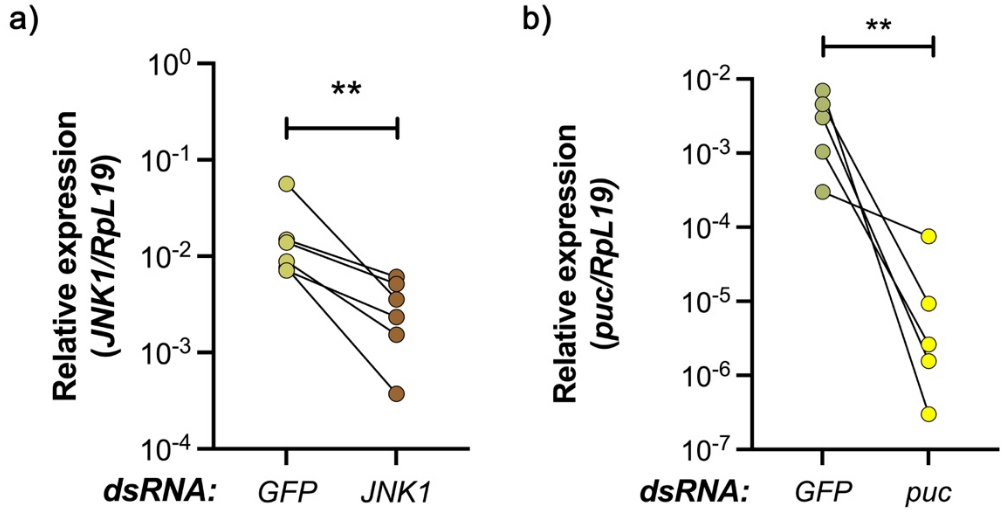


**Supplementary Figure 2. *dsJNK1* and *dspuc* knock down of gene expression.** Carcasses (whole body minus head, reproductive tract and gut) of unfed females from the groups used in Figure 1 (*dsJNK1*) and Figure 3 (*dspuc*) were dissected 48 hours after injection with the *dsRNA* indicated and subjected to qRT-PCR analysis using the delta delta CT method. Expression of *JNK1* (**a**) or *puc* (**b**) was calculated relative to that of the housekeeping gene, *RpL19,* in the same sample. Data show the results of 6 (*JNK1*) or 5 (*puc*) independent biological experiments. Connected dots represent data from the same experiment. Between-group differences in relative expression were compared using a Mann-Whitney test. *dsJNK1*, $p$=0.0022, Mann-Whitney U=0, median *dsGFP*=0.011, median *dsJNK1*=0.0030 Hodges-Lehmann estimate=-0.0081, 95%CI [-0.0035, -0.050]; *dspuc*, p=0.0079, Mann-Whitney U=0, median *dsGFP*=0.00303, median *dspuc*=0.0000026, Hodges-Lehmann estimate=-0.0030, 95%CI [-0.00029, -0.0070]. ** Denotes $p<0.01$. $p<0.05$ was taken to be significant.

Supplementary Figure 3

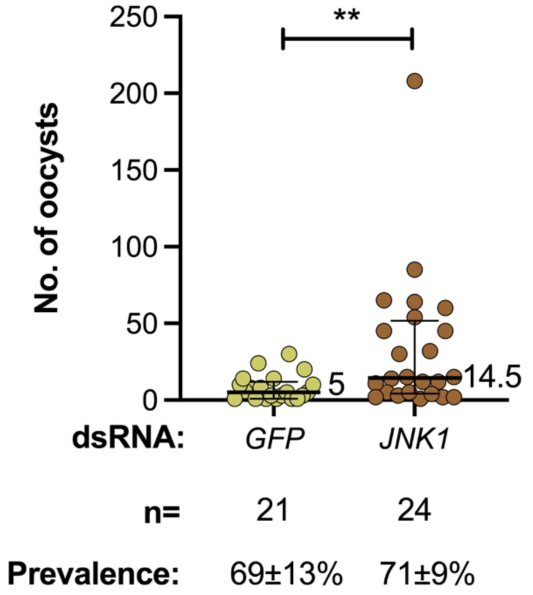


**Supplementary Figure 3. JNK depletion enhances low intensity *P. berghei* infection.** A subset of the experiments presented in Figure 1C in which infection intensity was low (median oocysts/ midgut ≤10). Data show the median and interquartile range of oocyst counts at d8 post infection in *dsGFP*- or *dsJNK1*-injected females from three independent biological replicates. Differences in infection intensity were compared using a Mann-Whitney test ($p$=0.0074, Mann-Whitney U=136, median *dsGFP*=5, median *dsJNK1*=14.5, Hodges-Lehmann estimate=10, 95%CI [1, 30]). ** denotes p<0.01; p<0.05 was taken to be significant. Females lacking oocysts at d8 post infection were not included in the analysis. The number of mosquitoes analyzed (n) and the mean ± SEM % infection prevalence are also shown. There was no significant difference in infection prevalence using Fisher's exact test on the absolute values ($p$=0.63).

### Supplementary Figure 4

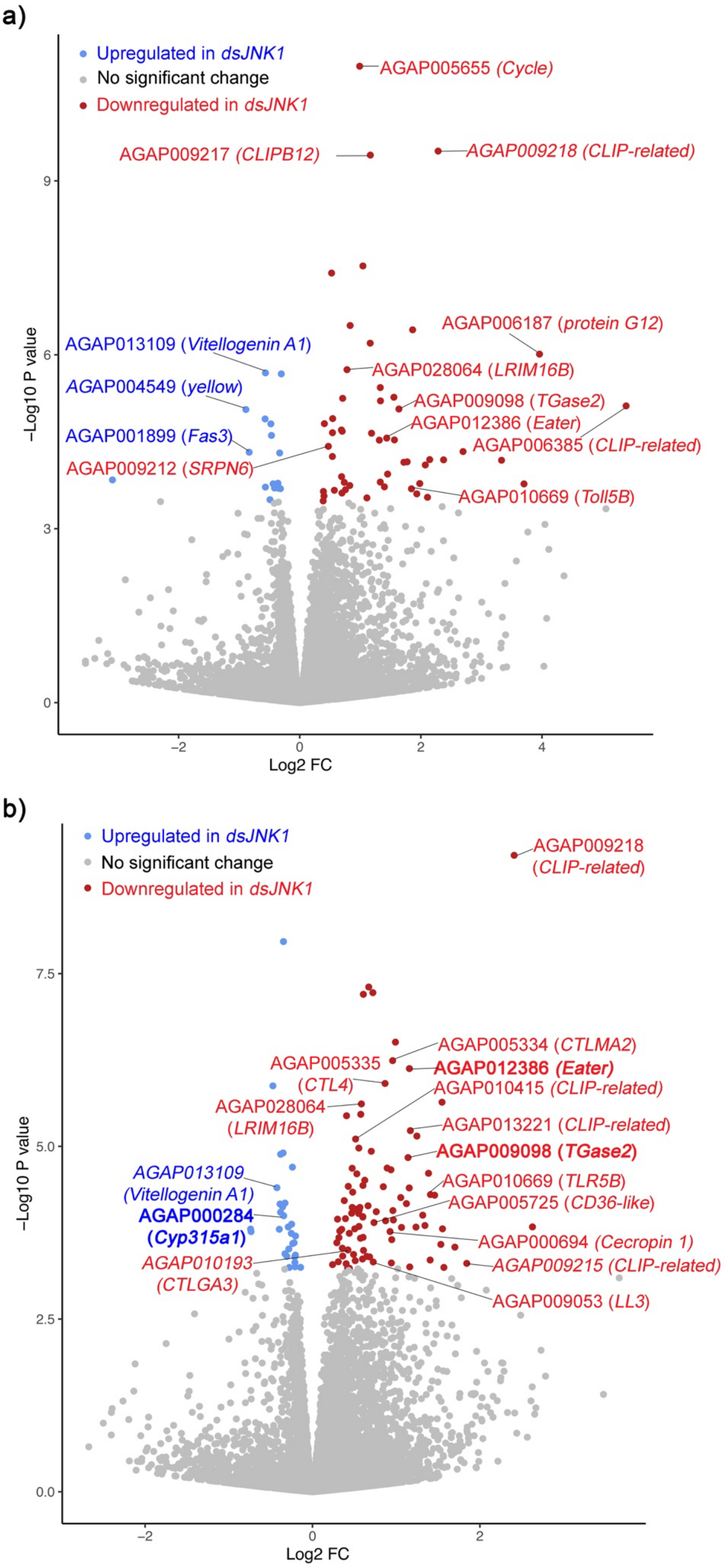

**Supplementary Figure 4. Volcano plots of *dsJNK1*-induced gene expression changes.** Plots compare RNA-seq gene expression analysis of *dsGFP*- (control) and *dsJNK1*-injected females 24hPBF. **a)** considers only those females fed on a *P. berghei*-infected mouse and represents four biological replicates (four samples per treatment). **b)** expression data from both infected and uninfected blood meals for each treatment (*dsGFP* vs *dsJNK1*), from the 4 biological replicates used in panel A, were pooled (eight samples per treatment). Genes whose expression was significantly altered by *dsJNK1* (FDR-adjusted $p<0.05$) are indicated in blue (upregulated by *dsJNK1*) or red (downregulated by *dsJNK1*). A number of genes of interest, including examples of the tendency towards immune function, are listed. In panel B, differentially expressed genes in bold text were verified by qRT analysis. Lists of all genes differentially expressed are presented in **Supplementary Tables 1 and 2**.

#### Supplementary Figure 5

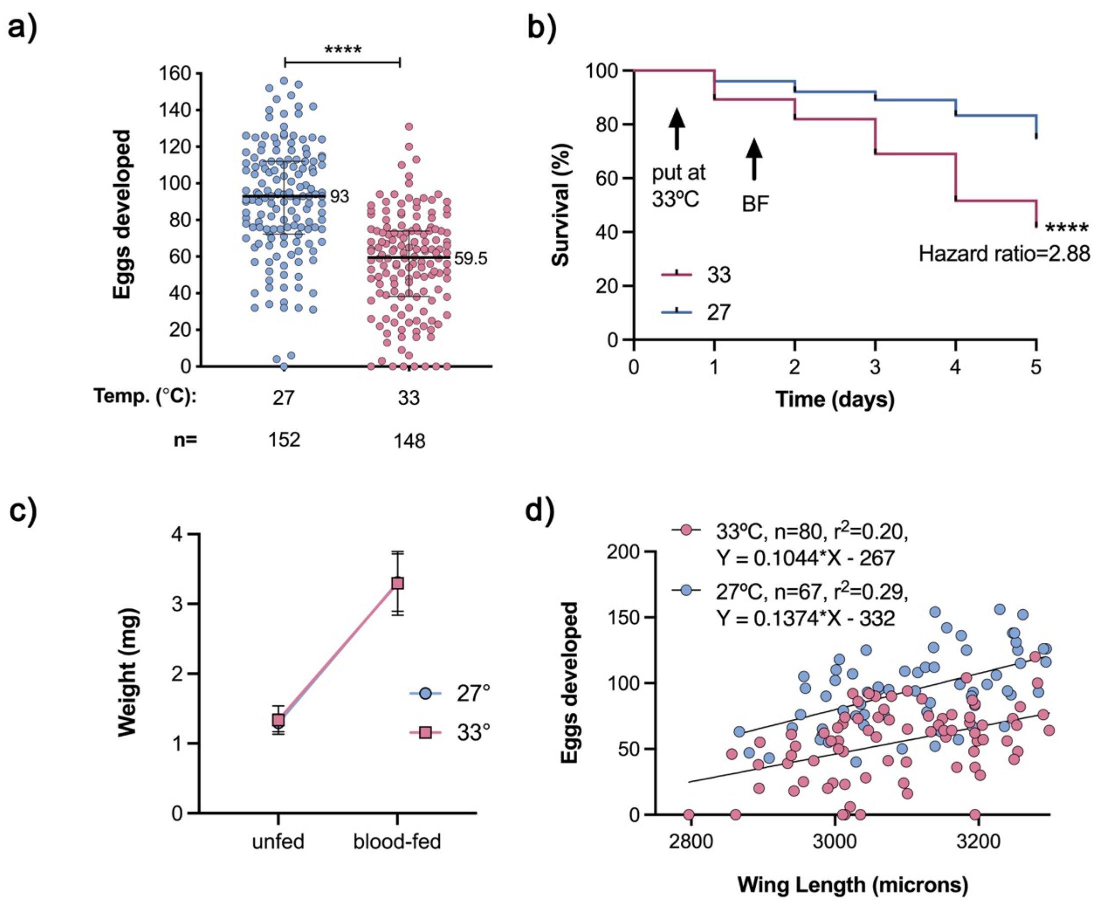


**Supplementary Figure 5. Heat stress reduces egg number and increases mortality without affecting blood meal size or mean size of survivors. a)** Virgin females were placed in an incubator at the temperature indicated then 24 hours later allowed to feed on the same blood. After a further 72 hours at either 27 or 33ºC the number of eggs developed was counted. The total number of mosquitoes analyzed (n) is shown. The data represent the median ± interquartile range of 6 independent biological replicates. Between group differences were analyzed using a Mann-Whitney test ($p<0.0001$, Mann-Whitney U=9.05, median 27ºC=93, median 33ºC=59, Hodges-Lehmann estimate=-34, 95%CI [-28, -41]). **b)** Daily mortality at 27ºC and 33ºC was monitored before and three days PBF and compared using a Log Rank Mantel-Cox test and used to calculate the hazard ratio (HR, the relative risk of death at 33ºC vs 27ºC). Death at 33ºC was more frequent than at 27ºC ($p<0.0001$, Chi square=180, df=1, HR=2.88, 95%CI [2.48, 3.35]. Data represent 5 independent biological replicates. **c)** The pre- and post-blood feeding weight of females exposed or not to 24-hour incubation at 33ºC. The data shown represent the mean ± SD of 3 independent experiments comprising a total of 59 unfed and 60 blood-fed controls and 151 unfed and 117 blood-fed heat-exposed females. **d)** Blood-fed females subjected or not to 33ºC had their eggs counted at day 3 PBF. At the same time wings were dissected and measured using ImageJ. The data show a Pearsons linear regression analysis of egg number and wing length for the two groups. The number of females analyzed (n), the value of $r^2$ and the equations of the lines of best fit are all indicated. While the slopes are indistinguishable (*p*=0.40) the difference in the y intercepts (27ºC, y intercept =-267.2, 95%CI [-415.5, -119.8]; 33ºC, y intercept=-332.4, 95%CI=-497.8, -167.1], $p<0.0001$) is highly significant. Data come from 4 independent biological replicates.

#### Supplementary Figure 6

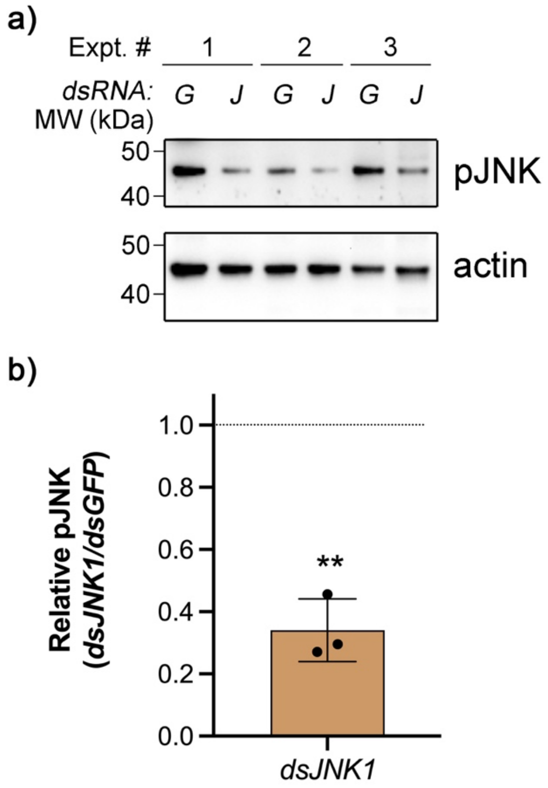


**Supplementary Figure 6. *dsJNK1* reduces pJNK levels after blood feeding under heat stress. a)** Virgin females were injected with *dsGFP* (*G*) or *dsJNK1* (*J*) then 24 hPBF under heat stress, were western blotted for pJNK and actin. Originals of the western blots presented here appear in full in Supplemantary Data **b)** Actin-normalized densitometry values for the pJNK signal were used to calculate the ratio of the signal in *dsJNK1*- and *dsGFP*-injected females. Deviation from a ratio of 1 (indicated by the dotted line) was analyzed using a one sample t-test (t(2)=11.33, p=0.0077, discrepancy=-0.659, 95%CI [-0.91, -0.41]). The graph shows the mean ± SEM of 3 independent biological replicates, each represented by a black circle. ** denotes *p*<0.01. *p*<0.05 was taken to be significant.

**Supplementary Table 1. Genes differentially expressed in reproductive tracts of *JNK1*-depleted relative to control females 24 hours after a *P. berghei*-infected blood meal.** Genes listed here passed an adjusted statistical significance cut-off of $p<0.05$ but no limit was placed on fold-change in expression. Genes highlighted in bright yellow have a literature-based connection to immunity. Those in pale yellow are over-expressed in hemocytes relative to other somatic cells [1]. Genes whose expression was increased by *dsJNK1* are highlighted in pink. Genes unique to this dataset are highlighted in blue while those common to Table S2 are highlighted in green.

**Supplementary Table 2. Genes differentially expressed in reproductive tracts of *JNK1*-depleted relative to control females 24 hours after blood feeding (*P. berghei*-infected or naïve blood meal).** Genes listed here passed an adjusted statistical significance cut-off of $p<0.05$ but no limit was placed on fold-change in expression. Genes highlighted in bright yellow have a literature-based connection to immunity. Those in pale yellow are over-expressed in hemocytes relative to other somatic cells [1]. Genes whose expression was increased by *dsJNK1* are highlighted in pink. Genes unique to this dataset are highlighted in purple while those common to Table S1 are highlighted in green.

Supplementary Table 1

| GeneID | AGAP | Base mean | log2(FC) | StdErr | Wald-Stats | P-value | P-adj |
|---|---|---|---|---|---|---|---|
| LOC1276339 | AGAP005655-PA. Protein cycle | 766.864410481227 | 0.989203212316402 | 0.145491746093843 | 6.79903320205094 | 1.05323544885637e-11 | 1.09083595438054e-07 |
| LOC1280175 | AGAP009217. Phenoloxidase-activating factor 3. CLIPB12 | 570.26228428483 | 1.16581966655812 | 0.185919410461239 | 6.27056456163395 | 3.59741347366184e-10 | 1.24194704489052e-06 |
| LOC4578285 | AGAP009218-PA. CLIP domain-containing serine protease HP8 | 116.557632389758 | 2.28503390996893 | 0.362963920088431 | 6.29548498763241 | 3.06440218785426e-10 | 1.24194704489052e-06 |
| LOC1276395 | AGAP005721. Carbohydrate sulfotransferase 11. chondroitin 4- | 665.752999660925 | 1.04317282726818 | 0.188124068607607 | 5.54513218318731 | 2.93732184849679e-08 | 7.60546059622032e-05 |
| LOC1279313 | AGAP009897-PA. proton-coupled amino acid transporter-like protein | 17832.1122045466 | 0.52578564128718 | 0.095673738228923 | 5.49561092751605 | 3.89360101159948e-08 | 8.06520513542717e-05 |
| LOC1277034 | AGAP006422-PA. myrosinase 1, cyanogenic beta-glucosidase | 1182.0592625908 | 0.832413093467567 | 0.162723066505032 | 5.11551995267879 | 3.12877869574792e-07 | 0.000540079349197686 |
| LOC1273384 | AGAP002587. Monocarboxylate transporter | 235.601604402882 | 1.86381787293411 | 0.366653645322869 | 5.08332017616479 | 3.70893712862061e-07 | 0.000548763740587481 |
| LOC1277346 | AGAP000834. UNC93-like protein | 346.094385128082 | 1.16331757438471 | 0.233512318051209 | 4.98182530195087 | 6.29872878092026e-07 | 0.000815449174799889 |
| LOC1276853 | AGAP006187. Protein G12. | 27.1756384511518 | 3.95859372833419 | 0.808407230241092 | 4.89678169646457 | 9.74189592237126e-07 | 0.00112107573408888 |
| LOC11175611 | AGAP013109-PA. Vitellogenin-A1 | 13582.2720193195 | -0.566241361842133 | 0.119254853104431 | -4.74816200013492 | 2.0527358842552e-06 | 0.0018432766302873 |
| LOC133392694 | AGAP028064. Toll-like receptor Tollo | 921.460388727335 | 0.780234521827038 | 0.16343053333545 | 4.7741049723283 | 1.80508216489242e-06 | 0.0018432766302873 |
| LOC3291767 | AGAP008638. Histone H1 | 14538.3704065156 | -0.304259214817393 | 0.064187788690741 | -4.74014171579147 | 2.13568789837285e-06 | 0.0018432766302873 |
| LOC1277224 | AGAP006638. Solute carrier organic anion transporter family member 74D | 239.511136890445 | 1.33007918074095 | 0.287344579021172 | 4.62886470756403 | 3.67675870995515e-06 | 0.00292924538146196 |
| LOC3291994 | AGAP009549-PA. Probable nuclear hormone receptor HR38. | 480.968901582461 | 0.708600663766522 | 0.156098372775205 | 4.5394493944339 | 5.64013142868256e-06 | 0.00389432274712435 |
| LOC1270615 | AGAP011322. Fibulin-1. | 312.008838238427 | 1.55456467375108 | 0.341772872822481 | 4.54853148792335 | 5.40215612728663e-06 | 0.00389432274712435 |
| LOC1276518 | AGAP005849-PA. Colmedin. | 104.924180917412 | 1.33325498176239 | 0.295084242101986 | 4.51821816124494 | 6.23622119977626e-06 | 0.00403678393538017 |
| LOC1276997 | AGAP006385-PA. Peptidase S1 family CLIP subfamily. | 13.9281322474652 | 5.38929517609251 | 1.20405670188446 | 4.47594799120155 | 7.60730031948884e-06 | 0.00463463584758506 |
| LOC1274690 | AGAP004549. Protein Yellow | 367.539050459084 | -0.889210969748771 | 0.200061393014759 | -4.44469048400144 | 8.80184928932923e-06 | 0.00479793437313594 |
| LOC1280050 | AGAP009098-PA. Tgase2. Protein-glutamine gamma-glutamyltransferase | 224.446254014907 | 1.63740545415659 | 0.368009012444099 | 4.44936237643178 | 8.61255962328706e-06 | 0.00479793437313594 |
| LOC1281412 | AGAP001763-PA. Long-chain fatty acid transport protein 4. Fatty acid | 4432.19376058224 | -0.572102976669074 | 0.131095825901095 | -4.3640060447134 | 1.2770204592528e-05 | 0.00629814328403867 |
| LOC1276450 | AGAP005781. 2-amino-3-ketobutyrate coenzyme A ligase, mitochondrial | 4289.35954163876 | 0.543835019390783 | 0.124537380142891 | 4.3668416564312 | 1.2605604270494e-05 | 0.00629814328403867 |
| LOC5667071 | AGAP006645. Insulin-like growth factor-binding protein complex acid labile | 16145.6109126193 | 0.406879087846775 | 0.094129104945850 | 4.3225640792063 | 1.54226269069356e-05 | 0.0069901888632647 |
| LOC1279952 | AGAP008990-PA. Phospholipase A2 group XV. lysophospholipase III | 5606.09371720924 | -0.476725670491861 | 0.110324319660931 | -4.32112948402512 | 1.55232542102045e-05 | 0.0069901888632647 |
| LOC1278211 | AGAP007753-PA. Carbohydrate (trehalose) transporter? | 2656.51358525931 | 0.688736791049172 | 0.161414371117355 | 4.26688643818729 | 1.98219995364249e-05 | 0.00855401871661468 |
| LOC1276633 | AGAP005962-PA. Leucnie rich repeat-containing receptor. | 1501.70912418605 | 0.700864543125879 | 0.164688937578808 | 4.25568683258094 | 2.08408248083935e-05 | 0.00863393690162127 |
| LOC1274497 | AGAP004350. Synaptic vesicle glycoprotein 2B | 459.713993564193 | 1.18551554549956 | 0.279721822003828 | 4.23819470718068 | 2.25324318036393e-05 | 0.0086432739329738 |
| LOC3291932 | AGAP009239-PA. RanBP-type and C3HC4-type zinc finger-containing | 1272.50035144606 | 0.540755594458903 | 0.127453328609404 | 4.24277341642535 | 2.20774312507532e-05 | 0.0086432739329738 |
| LOC1269735 | AGAP007491. Qsox1 Quiescin sulfhydryl oxidase 1. Protein disulphide | 7531.37559924021 | -0.466971549690573 | 0.110699501057736 | -4.21837086191582 | 2.46073840602694e-05 | 0.00910209559686464 |
| LOC133392670 | AGAP012386. Eater. EGF-like domain-containing protein. Tenascin-like | 247.044322971543 | 1.43598860738818 | 0.342358781294301 | 4.19439688960033 | 2.73598570095752e-05 | 0.00977124272579896 |
| LOC1276660 | AGAP005987. Protein anachronism. Secreted? | 3345.91777152587 | 1.31357740737837 | 0.314557286406773 | 4.17595606314999 | 2.96737095322473e-05 | 0.00991389063308017 |
| LOC1275497 | AGAP008648. Methyl transferase activity? | 239.538595390816 | 1.56334444756455 | 0.374190076058584 | 4.17794203425051 | 2.9415859474528e-05 | 0.00991389063308017 |
| LOC1280171 | AGAP009212-PA. SRPN6. Serine-type endopeptidase inhibitor activity. | 3628.8285380145 | 0.472468902459217 | 0.114687742209813 | 4.11961115770219 | 3.79512306727916e-05 | 0.0122831530024407 |
| LOC1281226 | AGAP001899. Fatty acid synthase 3 | 600.254810759349 | -0.83533332901996 | 0.205482304406055 | -4.0652324366056 | 4.79846155532524e-05 | 0.014616960684854 |
| LOC1277742 | AGAP008237. G-protein coupled receptor 143 | 21.015175622542 | 2.69653777152327 | 0.662275028964717 | 4.07162833202176 | 4.66856295250891e-05 | 0.014616960684854 |
| LOC1279813 | AGAP008843. Aquaporin FA-CHIP | 6817.15923226624 | -0.331525292354275 | 0.081688060763943 | -4.0584301947416 | 4.94036962018991e-05 | 0.0146192594732305 |
| LOC5667799 | AGAP011749-PA. Uncharacterised. Membrane protein, 2 MFS transporter | 958.531988796664 | 0.539120277991129 | 0.133916514294022 | 4.02579383755058 | 5.67834234274969e-05 | 0.0163362754566274 |
| LOC1281406 | AGAP001769-PA. Beat protein. Transmembrane protein. | 115.920274594887 | 2.14590131057977 | 0.536919913733478 | 3.99668787782189 | 6.42349047195382e-05 | 0.0175268257919254 |
| LOC4576502 | AGAP005889. Uncharacterized. Secreted? | 1279.93286898636 | 2.37484993800955 | 0.594551835372334 | 3.99435305169367 | 6.48711398119683e-05 | 0.0175268257919254 |
| LOC1278221 | AGAP007745. Secreted? Pertitrophic matrix associated protein | 27.1622071069829 | 3.33210108871982 | 0.83505681282336 | 3.99026873088295 | 6.59984750299403e-05 | 0.0175268257919254 |
| LOC1272376 | AGAP000785. Synaptic vesicle glycoprotein 2C | 154.74370499297 | 1.71933124741967 | 0.432900969664851 | 3.97165025698779 | 7.13764582350041e-05 | 0.0180303897058521 |
| LOC1277612 | AGAP008369-PA. Vitellogenin domain-containing protein | 145.452696531889 | 1.77402150861138 | 0.446236120160731 | 3.97552199040362 | 7.02250541509034e-05 | 0.0180303897058521 |
| LOC133393780 | AGAP028422. Uncharacterised. Secreted? | 50.747078390842 | 2.07348045281387 | 0.525477838425285 | 3.94589514760039 | 7.95024010916721e-05 | 0.019604913526344 |
| LOC3291074 | Non coding RNA | 186.89599404099 | 1.4508552301907 | 0.376029249068248 | 3.85835738519205 | 0.000114151645883143 | 0.0274946185212027 |
| LOC133393932 | AGAP011518. Phospholipid-transporting ATPase ABCA3-like. | 418.536812802376 | 0.691253693934814 | 0.180337218540396 | 3.83311719859965 | 0.000126529633448009 | 0.0297833503095688 |
| LOC1280041 | AGAP009087. PH domain leucine-rich repeat protein phosphatase | 38.093527902067 | -3.09234247373637 | 0.813521511961906 | -3.8011809500634 | 0.000144008083431755 | 0.0331442604467263 |
| LOC1271734 | AGAP000507. Zinc finger protein 567 | 3433.13681310597 | -0.359761582038328 | 0.095465431760069 | -3.76850107316863 | 0.000164230779346135 | 0.0342588154374289 |
| LOC1281521 | AGAP001652. Lipase 3 | 1092.24878441637 | 0.735577384257447 | 0.194766676806165 | 3.77671065872068 | 0.000158913158375824 | 0.0342588154374289 |
| LOC1270235 | AGAP006828. Cuticle protein 19. CPR60. | 68.2033911869979 | 1.33010179383985 | 0.351879885624443 | 3.77998813850774 | 0.0001568358301758 | 0.0342588154374289 |
| LOC1277620 | AGAP008359. Sodium-coupled monocarboxylate transporter 1 | 75.7740152603338 | 1.98125753656142 | 0.526158383036216 | 3.76551548058306 | 0.00016620582476417 | 0.0342588154374289 |
| LOC1271290 | AGAP009381. Clavesin-1. Cellular retinaldehyde-binding protein. | 2890.03682596124 | -0.439969882713171 | 0.116938107669602 | -3.76241664484828 | 0.000168279402324908 | 0.0342588154374289 |
| LOC133393409 | AGAP029067. Transmembrane protease serine 9-like. | 21.0847788003195 | 3.70267104375678 | 0.984282722940526 | 3.76179623746226 | 0.000168697459429263 | 0.0342588154374289 |
| LOC5667361 | AGAP004883. Secreted? Uncharacterised | 576.051698095248 | 0.826281632027858 | 0.220613130116566 | 3.74538737377632 | 0.00018011563807706 | 0.0358741858377714 |
| LOC11175555 | AGAP013003-PA. PH domain and PTB domain containing. | 2539.98891470737 | -0.566827929701019 | 0.151967267506086 | -3.72993434048762 | 0.000191529675421583 | 0.0359665192776066 |
| LOC1272074 | AGAP000190. Ptx1 pituitary homeobox homolog Ptx1. Transcription | 75.1557213766859 | 1.39836370236748 | 0.374638278335342 | 3.73257027707082 | 0.000189535791751396 | 0.0359665192776066 |
| LOC1273575 | AGAP002387. Acph-1 Acid phosphatase 1 | 15379.1179058466 | -0.318923464183258 | 0.085865684828961 | -3.71421324850004 | 0.000203836855938664 | 0.0359665192776066 |
| LOC1269756 | AGAP007460. Acyl-CoA-binding protein homolog | 2434.93998915707 | -0.362934748980966 | 0.097075353068859 | -3.73869100144837 | 0.000184980924149016 | 0.0359665192776066 |
| LOC1272472 | AGAP010669-PA. Protein toll. TOLL5B. Plasma membrane Toll-like | 29.0946216673559 | 1.84462933966205 | 0.49681469138389 | 3.7129122219067 | 0.000204887963442965 | 0.0359665192776066 |
| LOC3291198 | AGAP010938. Midnolin homolog. Homeodomain hit. Brinker DNA binding | 1546.39482514629 | -0.416313561038666 | 0.111888611676407 | -3.72078583156154 | 0.000198603815257012 | 0.0359665192776066 |
| SBT44_mgp08 | AGAP028371. COX3 cytochrome c oxidase subunit III | 83686.8143140048 | -0.35694206568068 | 0.095962937339743 | -3.71958253442137 | 0.000199552332727296 | 0.0359665192776066 |
| LOC1281766 | AGAP001410. Hemicentin-1. | 156.747576586242 | 0.75456541289867 | 0.203727010152822 | 3.70380644340015 | 0.00021238833622667 | 0.036661766638327 |
| LOC1270821 | AGAP011104-PA. Complement control module. | 565.09045120372 | 0.57118270347265 | 0.154457587203622 | 3.6979905863715 | 0.000217312946154039 | 0.0368968882511046 |
| LOC1269492 | AGAP003901. GAS2-like protein pickled eggs. | 882.00440106174 | 0.39106276778016 | 0.106097331460118 | 3.6858869341795 | 0.000227907524125805 | 0.0380715843124349 |
| LOC1270530 | AGAP001038-PA. cholesterol 7-desaturase nvd | 358.105179420181 | 0.696963297120828 | 0.189905901032153 | 3.67004549796915 | 0.000242507300629028 | 0.0398674303589658 |
| LOC4576344 | AGAP006367-PA. E3 ubiquitin-protein ligase TRAIP. Zinc/RING finger | 27.0025553157813 | 1.93299163069567 | 0.527878140084637 | 3.66181412699859 | 0.000250435539535155 | 0.0405275137963375 |
| LOC1275576 | AGAP008726. Eyes absent homolog 1 (eya) | 2709.76850480973 | 0.400783299639052 | 0.110112331123354 | 3.63976764046592 | 0.000272884174061238 | 0.0434809444731114 |
| LOC1276599 | AGAP005931. Differentially expressed in FDCP 8 homolog. | 70.9900134845315 | 2.1092459495745 | 0.581600825043406 | 3.62662131611846 | 0.000287153938466756 | 0.0450614142530332 |
| LOC1278354 | AGAP011505-PA. Constituent of cuticle | 277.665232634197 | 1.10898082730413 | 0.306325060611347 | 3.62027457071545 | 0.0002942905560136 | 0.0454920490840725 |
| LOC1276231 | AGAP005547. Protein lethal(2)essential for life. | 7082.1115562169 | -0.491570407663945 | 0.136467706029957 | -3.60210061387004 | 0.000315656138611439 | 0.0480772151117452 |
| LOC1272816 | AGAP003441. Anosmin-1. | 2548.45326360127 | 0.38665820083191 | 0.107727068660389 | 3.58923904307519 | 0.000331644631088035 | 0.049780339770707 |

Immune connection: 16/70 (22.9%)

JNK has suppressive effect; 17/70 (24.3%)

Over represented in hemocytes: 13/70 (18.6%)

Total 29/70 (41.4%)

Unique to this dataset

Shared with all dsGFP vs dsJNK dataset

29 OF 69 signficant differences also observed in dsGFP vs dsJNK global comaprison; 40 unique to IBF only data set.

Supplementary Table 2

| GeneID | AGAP | Base mean | log2(FC) | StdErr | Wald-Stats | P-value | P-adj |
|---|---|---|---|---|---|---|---|
| LOC4578285 | AGAP009218-PA. CLIP domain-containing serine protease HP8 | 705.391.000.077.941 | 2.40946962685715 | 0.389443836650801 | 6.1869502097619 | 6.13393283584217e-10 | 6.20079270375285e-06 |
| LOC1271734 | AGAP000507-PA. Zinc finger protein 567 | 3241.64631654535 | -0.346620808933571 | 0.0606333066045956 | -5.71667336558055 | 1.08629637085557e-08 | 5.49068500648946e-05 |
| LOC1276339 | AGAP005655-PA. Protein cycle | 740.419228910626 | 0.721347727268122 | 0.133093165934098 | 5.4198705260742 | 5.96422142201652e-08 | 0.000127350272567014 |
| LOC1277034 | AGAP006422-PA. Myrosinase 1, cyanogenic beta-glucosidase | 991.290839047236 | 0.672119191000686 | 0.123244745238404 | 5.45353223539504 | 4.93789896474482e-08 | 0.000127350272567014 |
| LOC3291994 | AGAP009549-PA. Probable nuclear hormone receptor HR38. | 421.456004608192 | 0.607912561984743 | 0.11236617455675 | 5.41010285686749 | 6.29885609689453e-08 | 0.000127350272567014 |
| LOC4577966 | Non coding RNA | 369.865893600866 | 0.99137395770654 | 0.193734389541076 | 5.11718110581677 | 3.1013580055091e-07 | 0.000522527134628192 |
| LOC1276042 | AGAP005334-PA. CTLMA2. C-type lectin 37Db | 247.567056786962 | 0.957822565748737 | 0.19158379456208 | 4.99949678905836 | 5.74801294673631e-07 | 0.000830095183979391 |
| LOC133392670 | AGAP012386. Eater. EGF-like domain-containing protein. tenascin-like | 195.93270906853 | 1.15796853708362 | 0.234012803950184 | 4.94831273134151 | 7.48595651591826e-07 | 0.000945944180242721 |
| LOC11175555 | AGAP028653. (A0A1S4HF38 ) PH domain-containing protein, phosphotyrosine-binding. | 2416.88951981679 | -0.473009072947943 | 0.0978309149804981 | -4.83496523611411 | 1.33168905665971e-06 | 0.0013462044673773 |
| LOC1276043 | AGAP005335-PA. C-type lectin (CTL). CTL4 | 423.887122428188 | 0.867080741216475 | 0.178757091049236 | 4.85060892480988 | 1.23083017361561e-06 | 0.0013462044673773 |
| LOC4576344 | AGAP006367-PA. E3 ubiquitin-protein ligase TRAIP | 23.062191335664 | 1.54769885657375 | 0.327531668051957 | 4.72534111214015 | 2.29729398868969e-06 | 0.00204766433442006 |
| LOC133392694 | AGAP028064. Toll-like receptor Tollo. Leucine-rich immune protein (TM) | 807.862289366705 | 0.584383176573161 | 0.123971319073465 | 4.71385785793615 | 2.43070254357906e-06 | 0.00204766433442006 |
| LOC133393932 | AGAP011518. Phospholipid-transporting ATPase ABCA3-like | 384.809082370928 | 0.576844324151129 | 0.124250480791136 | 4.64259229001135 | 3.44064869686911e-06 | 0.00260392127346134 |
| LOC5667799 | AGAP011749-PA. Major facilitator superfamily associated domain-containing protein | 854.835286413063 | 0.407176467758936 | 0.0878884629998655 | 4.63287732952571 | 3.60618239474318e-06 | 0.00260392127346134 |
| LOC1271098 | AGAP010858-PA. Sodium-dependent nutrient amino acid transporter 5 | 178.284581265777 | 1.16931514382752 | 0.258184388227164 | 4.52899244550253 | 5.92656176982817e-06 | 0.00399410752874619 |
| LOC11175927 | AGAP013221-PA. Fibronectin type 1 (CLIP family serine protease, snake) | 136.872243284992 | 1.24727835396482 | 0.277743056264649 | 4.49076340823566 | 7.09683482215203e-06 | 0.00448386895107093 |
| LOC1272633 | AGAP010415-PA. CLIP family S1 Peptidase | 258.834172862801 | 0.514868531310043 | 0.115195974227529 | 4.46950108076782 | 7.84022630492888e-06 | 0.00466216751273683 |
| LOC1271001 | AGAP010964-PA. Ig-like domain-containing | 495.288629144486 | 0.553840894347178 | 0.125737844285344 | 4.40472713282976 | 1.05917113956743e-05 | 0.00594842280549285 |
| LOC1275277 | AGAP010546-PA. Putative CLIP family serine protease | 397.378195368638 | 0.702559523879058 | 0.160349456110458 | 4.3814275453176 | 1.17904252612311e-05 | 0.00627312678767291 |
| LOC1280232 | AGAP009268-PA. BicC protein bicaudal C, Ets-like Transcription factor? | 13020.9937696459 | -0.380630897461822 | 0.0873094547920937 | -4.35956103915896 | 1.30323599589805e-05 | 0.00627352984882543 |
| LOC1271249 | AGAP009338-PA. Cell division cycle 20 protein fzy | 1576.17488057656 | -0.351035545734095 | 0.0803417541752868 | -4.36927907957072 | 1.24657376325466e-05 | 0.00627352984882543 |
| LOC1280050 | AGAP009098-PA. Tgase2. Protein-glutamine gamma-glutamyltransferase E, annulin | 168.45715943839 | 1.14162714937758 | 0.263318258404286 | 4.3355411671635 | 1.45401958037059e-05 | 0.00668121997180288 |
| LOC1276486 | AGAP005817-PA. Cyclin-dependent kinase 4 | 1467.99670452627 | -0.239731663374595 | 0.0562122050601868 | -4.26476177402955 | 2.00115600932046e-05 | 0.00839569912492047 |
| LOC1276518 | AGAP005849-PA. Transmembrane protein. 2 Ig-like domains. Signaling Receptor? | 89.0445726797613 | 0.89227697363752 | 0.209559749576129 | 4.25786428664047 | 2.06389172750766e-05 | 0.00839569912492047 |
| LOC5666864 | AGAP006380-PA. phospholipid-transporting ATPase ABCA3. | 1220.13704604241 | 0.47423438496558 | 0.111413522462085 | 4.25652447284363 | 2.07629318550808e-05 | 0.00839569912492047 |
| LOC1279406 | AGAP009985-PA. 4-nitrophenylphosphatase-related | 101.048086513619 | 0.937277541879503 | 0.220782230384592 | 4.24525805472121 | 2.18341992087108e-05 | 0.00848930460772529 |
| LOC1270530 | AGAP001038-PA. cholesterol 7-desaturase nvd, cholesterol 7-dehydrogenase | 333.842385620377 | 0.529735397748947 | 0.125677740562502 | 4.21502960968255 | 2.49745420439263e-05 | 0.00901670162578755 |
| LOC1272689 | AGAP010350-PA. Contains TRAF-like domain | 62.3739192312009 | 1.38635648907314 | 0.328603307583902 | 4.21893650208972 | 2.45457383992161e-05 | 0.00901670162578755 |
| LOC1275644 | AGAP008783. Arginase. | 418.239988726305 | 0.624645542388166 | 0.149922890281862 | 4.16644543881061 | 3.0938589717894e-05 | 0.0107847656364893 |
| LOC1281525 | AGAP001648-PA. Serine protease grass, CLIPB17 | 185.798148758518 | 0.846342144403553 | 0.205584159927345 | 4.11676728743429 | 3.84223678233532e-05 | 0.0118763744344255 |
| LOC1273328 | ncRNA. miRNA? | 889.407300784346 | 0.603254622464294 | 0.14619146824413 | 4.1264694151433 | 3.68374969637801e-05 | 0.0118763744344255 |
| LOC11175611 | AGAP013109. Vitellogenin-A1 lipid transporter. | 11375.6654812064 | -0.42478528818636 | 0.103366138172662 | -4.10952073566688 | 3.96481114014615e-05 | 0.0118763744344255 |
| LOC1276450 | AGAP005781. 2-amino-3-ketobutyrate coenzyme A ligase, mitochondrial | 3976.26470716798 | 0.427587777674811 | 0.103794270813316 | 4.11957012968346 | 3.79579885283086e-05 | 0.0118763744344255 |
| LOC1280199 | AGAP009246-PA. Cyp4C27. cytochrome P450 4C1. Linked to heat stress resistance in | 84.9149135320969 | 1.15592194014724 | 0.281396723054035 | 4.10780170999105 | 3.99442804204635e-05 | 0.0118763744344255 |
| LOC4577218 | AGAP001496-PA. Zinc finger C2H2 domain containing. Regulation of gene expression. | 610.490271978203 | 0.480375807116642 | 0.117882976851521 | 4.07502270426794 | 4.60098579307625e-05 | 0.0132889615377737 |
| LOC133393780 | AGAP028422. Uncharacterised, Protein coding | 37.6167193199875 | 1.45955745797533 | 0.360270076183215 | 4.05128694960797 | 5.09366936115442e-05 | 0.0139167306951108 |
| LOC1272472 | AGAP010669-PA. protein Toll-like receptor 5B | 21.3655775331987 | 1.40930059148857 | 0.347487952610422 | 4.05568187587953 | 4.99882574928017e-05 | 0.0139167306951108 |
| LOC11175747 | AGAP013185-PA. Signal peptide, probably secreted. | 79.9951222232689 | 1.05407444685457 | 0.261387126566222 | 4.03261805851604 | 5.51588887997627e-05 | 0.0146737159704421 |
| LOC5667351 | AGAP006402-PA. Protein suppressor 2 of zeste. | 447.35551010363 | 0.379720420071215 | 0.0947246851693446 | 4.00867439561681 | 6.10605378212827e-05 | 0.0158272045342397 |
| LOC3291761 | AGAP008584-PA. Lysosomal alpha-mannosidase. Glycoside hydrolase 38 family. | 2253.1690879283 | -0.387820452141353 | 0.0973932613797837 | -3.98200498316873 | 6.83363467013506e-05 | 0.0164479078286655 |
| LOC1279952 | AGAP008990-PA. Phospholipase A2 group XV. lysophospholipase III | 5218.77585978466 | -0.328443941562487 | 0.082327205597262 | -3.98949459270132 | 6.62142285937808e-05 | 0.0164479078286655 |
| LOC4397705 | AGAP012647-PA. Luciferin sulfotransferase | 32.7104936715948 | 1.12229342510927 | 0.281632189286288 | 3.98496147742695 | 6.74910738371172e-05 | 0.0164479078286655 |
| LOC1271942 | AGAP000321-PA. Uncharacterized. | 895.184978515152 | 0.474673987765232 | 0.120021429411691 | 3.95491030303457 | 7.65633709647893e-05 | 0.0168342960011186 |
| LOC1281766 | AGAP001410-PA. Hemicentin-1. | 140.992336301595 | 0.566562783855145 | 0.143259991004991 | 3.95478723599394 | 7.66027911812696e-05 | 0.0168342960011186 |
| LOC1276621 | AGAP005952-PA. Basic proline-rich protein | 316.168000748944 | 0.663662455514838 | 0.167255645151684 | 3.96795250117246 | 7.2492777541128e-05 | 0.0168342960011186 |
| LOC4577473 | AGAP012327-PA. CLIP domain-containing serine protease B4-like | 318.340133464818 | -0.357381810272575 | 0.0902721623509787 | -3.95893707390183 | 7.52840671030211e-05 | 0.0168342960011186 |
| LOC1281521 | AGAP001652. Lipase 3. Lysosomal acid lipase-like | 914.235588907869 | 0.513812886605444 | 0.130360114783225 | 3.94148844882393 | 8.09775353879595e-05 | 0.0174170618135507 |
| LOC1272903 | AGAP003049-PA. Stearoyl-CoA desaturase 5. Unsaturated fatty acid biosynthesis | 498.550407101037 | 0.558014840283688 | 0.141815714359939 | 3.93478848801906 | 8.32699953074857e-05 | 0.0175370079700703 |
| LOC1277224 | AGAP006638-PA. Solute carrier organic anion transporter family member 74D. Oatp58Dc | 219.661318156882 | 0.960864989835863 | 0.244514900394106 | 3.92967867515294 | 8.50594518615751e-05 | 0.0175482856911972 |
| LOC11175545 | AGAP013260-PA. Endochitinase. chitinase 5-5 | 197.603171338519 | 0.761735568447805 | 0.194287661665518 | 3.92065848092399 | 8.83073382784959e-05 | 0.0178539776531463 |
| LOC1281412 | AGAP001763-PA. Long-chain fatty acid transport protein 4. Fatty acid transporter protein 1 | 4172.28654501035 | -0.381275728584115 | 0.0973687366907698 | -3.91579208627299 | 9.01078785069948e-05 | 0.0178607949770041 |
| LOC11175456 | AGAP013533-PA. Juvenile hormone binding protein 1. | 590.114002361849 | 0.551892152629015 | 0.141288553359555 | 3.90613492392795 | 9.37841591977293e-05 | 0.0178880012326386 |
| LOC1275273 | AGAP010543-PA. Protein yellow | 336.327256051624 | 0.468379202477191 | 0.119849162679163 | 3.90807238037236 | 9.30354370398877e-05 | 0.0178880012326386 |
| LOC1280320 | AGAP012394-PA. Peptide methionine sulfoxide reductase | 84.5653585166257 | 1.31735532633431 | 0.338419204727094 | 3.89267307508935 | 9.9145719764222e-05 | 0.0185604459462319 |
| LOC1271975 | AGAP000284-PA. Cytochrome P450 family protein sad. | 507.736694955688 | -0.343504116484226 | 0.0883505026227344 | -3.88797014490142 | 0.00010108605662258 | 0.0185796172072316 |
| LOC1269400 | AGAP002156-PA. adipokinetic hormone/corazonin-related peptide receptor variant I. | 172.070637252482 | 0.601698380426883 | 0.155047631068527 | 3.8807324967187 | 0.00010414233634971 | 0.0187443270888247 |
| LOC1277021 | AGAP006410-PA. One cut domain family member 3. Secreted? | 181.097335099555 | 0.601276662885325 | 0.155082441599092 | 3.87714209736072 | 0.00010569063646879 | 0.0187443270888247 |
| LOC1277615 | AGAP008364-PA. Thioester-containing protein 1 allele R1-like. | 2421.26963950843 | 0.394076642792792 | 0.101972124938403 | 3.86455262191346 | 0.00011129299576627 | 0.0193976016241602 |
| LOC1280951 | AGAP011685. Protein unzipped. Secreted? | 771.241693501693 | 0.299324217201629 | 0.0775383703688379 | 3.86033670526979 | 0.00011323088989733 | 0.0194008655249511 |
| LOC1274755 | AGAP005072. Basement membrane-specific heparan sulfate proteoglycan core protein. | 82.4941829796901 | 0.966603473114255 | 0.250835602971293 | 3.85353379529971 | 0.00011642518480826 | 0.0196157032204463 |
| LOC1280013 | AGAP009056-PA Unchacracterized (also listed as AGAP028153). | 54.3894515854909 | 0.873378229879229 | 0.227032011432126 | 3.84693869542857 | 0.00011960288058937 | 0.0198207462275087 |
| LOC1276398 | AGAP005725-PA. Class B Scavenger Receptor (CD36 domain). | 260.394210575052 | 0.73484907771254 | 0.191755800192894 | 3.8322130385278 | 0.00012699569133734 | 0.020706442640794 |
| LOC1269746 | AGAP007477-PA. Replication factor C subunit RfC4 | 1619.55900546065 | -0.249141115416453 | 0.0652336704194234 | -3.81921044476857 | 0.00013387951095612 | 0.0214823488294524 |
| LOC1272150 | AGAP000117. Shifted protein. | 405.68283978772 | 0.349082154407186 | 0.0924307428544144 | 3.77668883346548 | 0.00015892707802198 | 0.0217107274557326 |
| LOC1273921 | AGAP003248. CLIP domain-containing serine protease B4-like | 95.2501783151417 | 1.23628604025948 | 0.32576567152885 | 3.79501632095692 | 0.00014763374413960 | 0.0217107274557326 |
| LOC1272847 | AGAP003475. Putative aminopeptidase W07G4.4. leucyl aminopeptidase | 5800.00156835073 | -0.289754094219707 | 0.0763278562901836 | -3.79617754647948 | 0.00014694427781941 | 0.0217107274557326 |
| LOC11175584 | AGAP012974-PA. Haparanase-Like Protein | 1893.0770115336 | -0.396401047751143 | 0.104944203272065 | -3.77725529749828 | 0.00015856617160202 | 0.0217107274557326 |
| LOC133391818 | ncRNA. miRNA? | 265.035742701701 | -0.739193198034359 | 0.195612177295666 | -3.77887107159527 | 0.00015754096022183 | 0.0217107274557326 |
| LOC4577091 | AGAP004316-PA. Uncharacterised. Secreted? | 54.394756379182 | 1.55360521286269 | 0.410902018826075 | 3.78096271539688 | 0.00015622307183007 | 0.0217107274557326 |
| LOC1270324 | AGAP006736. Zinc finger protein GLIS2. | 84.4853296718713 | 1.34331424733676 | 0.352744998434708 | 3.80817376092548 | 0.00013999688886398 | 0.0217107274557326 |
| LOC1270076 | AGAP007029. UDP-glycosyltransferase UGT5. | 257.164561890556 | 0.555697648057968 | 0.146206275216096 | 3.80077836766334 | 0.00014424227527681 | 0.0217107274557326 |
| LOC1278221 | AGAP007745. Uncharacterised. Secreted? | 18.5629313478975 | 2.6282530844427 | 0.69223798585823 | 3.79674784992363 | 0.00014660677691972 | 0.0217107274557326 |
| LOC1279487 | AGAP010057. Neuronal acetylcholine receptor subunit non-alpha-3 | 154.760530368675 | 0.504802232685245 | 0.133524708043545 | 3.78059042466223 | 0.00015645688055369 | 0.0217107274557326 |
| LOC1280823 | AGAP011824. Peroxiredoxin-6, TPX4. Thioredoxin-dependent peroxidase. | 48.8995796789392 | 1.06377523673283 | 0.280494574795057 | 3.79249843783994 | 0.00014913918901494 | 0.0217107274557326 |
| LOC1272282 | AGAP000694. Cecropin 1 | 204.194463280421 | 0.92713621799613 | 0.24669421647852 | 3.75824059124976 | 0.00017111231246678 | 0.0225337848287718 |
| LOC1274676 | AGAP004531. Cathepsin B | 11106.6252703985 | -0.733768015975461 | 0.195282418112308 | -3.75747096471054 | 0.00017163927508313 | 0.0225337848287718 |
| LOC1279313 | AGAP009897. Proton-coupled amino acid transporter-like protein pathetic | 16728.4060061777 | 0.326321057228586 | 0.0867310765728807 | 3.76244675061051 | 0.00016825914058495 | 0.0225337848287718 |
| LOC1281270 | AGAP001854-PA. Ogg1 8-oxoguanine DNA glycosylase. Involved_in base-excision repair | 602.373438633384 | -0.252476356970588 | 0.0674590588273557 | -3.74266053157867 | 0.00018208213650978 | 0.02329959896174 |

| | | | | | | | |
|---|---|---|---|---|---|---|---|
| LOC1279152 | AGAP009745. Facilitated trehalose transporter Tret1. | 842.619953275499 | 0.43957570740732 | 0.117422688739795 | 3.74353297582384 | 0.00018145077637593 | 0.02329959896174 |
| LOC1276587 | AGAP005920. Histone chaperone (asf1). | 2363.25834206941 | -0.202873009243653 | 0.0545194283197666 | -3.72111402294545 | 0.00019834584953151 | 0.0250634774114258 |
| LOC1273920 | AGAP003246. CLIP domain-containing serine protease B4-like | 353.80880295422 | 0.604583012720485 | 0.16298752525643 | 3.70938212460917 | 0.00020776565756418 | 0.0258222957925556 |
| LOC1280786 | AGAP011871. E3 ubiquitin-protein ligase. Involved_in proteasome-mediated ubiquitin- | 985.359905513579 | 0.316195683373514 | 0.0852894330463012 | 3.70732542215236 | 0.00020945971460971 | 0.0258222957925556 |
| LOC4578297 | AGAP010047-PA, putative svwc domain-containing protein. SVWC domain containing | 354.16880565076 | 0.573731328808951 | 0.154977950829812 | 3.70201906617664 | 0.00021389054041846 | 0.0260508370251844 |
| LOC1272074 | AGAP000190. Ptx1 pituitary homeobox homolog Ptx1. Homeobox domain-containing | 67.0434241869989 | 0.947073445419257 | 0.256649816773637 | 3.69013879427223 | 0.00022413173073176 | 0.0269731864996121 |
| LOC1269492 | AGAP003901. GAS2-like protein pickled eggs. | 809.144386917076 | 0.29071713977848 | 0.0793446397716943 | 3.66397957839354 | 0.00024832659308122 | 0.0292400344345176 |
| LOC3291494 | AGAP012040. Mucin-5AC. | 5181.89479651138 | -0.219557927372917 | 0.0599305314094109 | -3.66354047276876 | 0.00024875288963977 | 0.0292400344345176 |
| LOC1276468 | AGAP005800. Mcm7 minichromosome maintenance 7. DNA replication licensing factor | 5596.59467134594 | -0.235405279133434 | 0.0643724889894956 | -3.65692367700507 | 0.00025526034403673 | 0.0296600783662909 |
| LOC1276599 | AGAP005931. Relative of mammalian protein Differentially expressed in FDCP 8 homolog | 63.300526908524 | 1.5334793513331 | 0.420190919030967 | 3.64948237070323 | 0.00026276927160848 | 0.0301856200760249 |
| LOC1277612 | AGAP008369-PA. Vitellogenin domain-containing protein | 118.841946463281 | 1.70113665833743 | 0.469119806085076 | 3.6262307331124 | 0.00028758840619029 | 0.0326655190806479 |
| LOC3291932 | AGAP009239-PA. RanBP-type and C3HC4-type zinc finger-containing protein 1. | 1152.6818141719 | 0.356369313383649 | 0.0985244466404825 | 3.61706485583265 | 0.00029796266437644 | 0.0334678286020161 |
| LOC1280763 | AGAP011897-PA. THROMBOSPONDIN-TYPE LAMININ G DOMAIN AND EAR REPEAT- | 12280.3256357226 | -0.282860397983254 | 0.0783545815265462 | -3.6100045775552 | 0.00030619160151875 | 0.0340141857115725 |
| LOC1279613 | AGAP010193. CTLGA3. | 881.624492555049 | 0.423968462555216 | 0.117704335864183 | 3.60197829113472 | 0.00031580474226945 | 0.0347007623869775 |
| LOC3290136 | AGAP006257. Mucin-2 | 128.961311088439 | 0.6097557252566 | 0.169474538440095 | 3.59791937401933 | 0.00032077301656270 | 0.0348676819831443 |
| LOC4576148 | AGAP000575. Chromosome-associated kinesin KIF4A | 1193.36744503633 | -0.328392757694985 | 0.0920217296366562 | -3.56864361267311 | 0.00035883414366397 | 0.0385899399819052 |
| LOC1276917 | AGAP006263. ARR2. Arrestin homolog. | 485.326620246461 | 0.49069303673785 | 0.137733613864807 | 3.56262369779603 | 0.00036716680631612 | 0.0390704131057865 |
| LOC1280518 | AGAP012167. Glucose-6-phosphate isomerase. | 6224.23551014408 | -0.20431881497467 | 0.0574493825163533 | -3.55650149793184 | 0.00037582633845087 | 0.03957529641041 5 |
| LOC1269857 | AGAP007312-PA. Uncharacterised. | 227.383034942501 | 0.64925216471586 | 0.183125058858464 | 3.54540317290854 | 0.00039201298249094 | 0.0398486389237462 |
| LOC1269609 | AGAP007619. Cndp2 Cytosolic non-specific dipeptidase 2 | 7101.67348449148 | -0.212224099457694 | 0.0599158324940232 | -3.54203706472515 | 0.00039704967932933 | 0.0398486389237462 |
| LOC3291696 | AGAP008299-PA. Uncharacterised | 1170.08925568124 | -0.306776455805283 | 0.0865090936455129 | -3.54617581664139 | 0.00039086533358736 | 0.0398486389237462 |
| LOC133393029 | AGAP028641. Peptidase S1 domain-containing protein. CLIP subfamily. | 109.205050167285 | 0.682428355096042 | 0.192704559377163 | 3.54131919505022 | 0.00039813161848831 | 0.0398486389237462 |
| LOC1279404 | AGAP009982. Putative uncharacterized protein DDB_G0290521. Secreted? | 1079.26005231409 | 0.361097699670224 | 0.101753078771029 | 3.54876436203752 | 0.00038704326711359 | 0.0398486389237462 |
| LOC1277575 | AGAP008404-PA. UDPGT domain-containing protein. 563. | 294.876849703244 | 0.597079859304701 | 0.169511456015692 | 3.52235697420609 | 0.00042772776930555 | 0.0423911766657833 |
| LOC1270234 | AGAP006829. Cpr62Bc Cuticular protein 62Bc, cuticle protein 8, cuticular protein RR-1 | 35.1989921031188 | 1.40580905428425 | 0.400116908696625 | 3.51349574019168 | 0.00044225150493859 | 0.043405053042954 |
| LOC1269987 | AGAP007119. Sideroflexin-2. Mitochondrial inner membrane mono ion transporter | 399.22070686428 | 0.510607659139422 | 0.145571021612207 | 3.50761884806752 | 0.00045213634262770 | 0.0439485219963792 |
| LOC1276784 | AGAP006115-PA. Anion exchange protein 2 | 811.720913317738 | 0.308815062548158 | 0.088281320785519 | 3.49807931961541 | 0.00046862173804200 | 0.0449070862931553 |
| LOC1280011 | AGAP009053. LL3. Lipopolysaccharide-induced tumor necrosis factor-alpha factor | 129.274878701675 | 0.725441545079402 | 0.207458951032786 | 3.4967955900093 | 0.00047088249550642 | 0.0449070862931553 |
| LOC1280005 | AGAP009047. RfC38 replication factor C subunit RfC38 | 1482.76257797104 | -0.209902964841879 | 0.0601011697932128 | -3.4924938327171 | 0.00047853264626154 | 0.0452101544024105 |
| LOC1269855 | AGAP007314. Membrane protein. | 164.890589250501 | 0.942275216673222 | 0.270193360071689 | 3.48741070625574 | 0.00048772175599346 | 0.045651659549425 |
| LOC4578288 | AGAP009215. CLIP domain-containing serine protease HP8. CLIPB18. | 18.9896845816042 | 1.84273722646401 | 0.52897059940391 | 3.48362882273716 | 0.00049466499739991 | 0.0458767748506027 |
| LOC1281338 | AGAP002085-PA. Secreted, uncharactacterised. | 1169.38823202363 | 0.402230657177023 | 0.115553894969002 | 3.48089224759515 | 0.00049974649169586 | 0.0459267025868495 |
| LOC1281759 | AGAP001417. Mpc1 mitochondrial pyruvate carrier. | 2610.82231493382 | 0.240113905146643 | 0.0691442247804479 | 3.47265307997987 | 0.00051534102668488 | 0.0469331751239417 |
| LOC1273406 | AGAP002559. Alpha-tocopherol transfer protein-like. | 25.5700299130463 | 1.16171244104277 | 0.336555594757348 | 3.45176980902769 | 0.00055692263336781 | 0.0494225448269321 |
| LOC1273879 | AGAP003205. Monocarboxylate transporter 3 | 6725.19681121263 | -0.276734488435932 | 0.0802857175983165 | -3.44687071018637 | 0.00056711991294135 | 0.0494225448269321 |
| LOC1277279 | AGAP006690. BTB/POZ domain-containing protein 9. Galactose-binding domain-like 1 hit. | 30.1243600705941 | 1.56990654662871 | 0.455374571109456 | 3.44750595713733 | 0.00056578792975535 | 0.0494225448269321 |
| LOC1278873 | AGAP010789. Multiple epidermal growth factor-like domains protein 10. | 20405.9023836734 | -0.141850581028657 | 0.0411405011213011 | -3.44795462287678 | 0.00056484892541993 | 0.0494225448269321 |
| LOC1280807 | AGAP011845. Histone H2A family | 21621.0325847602 | -0.207626572092585 | 0.0601309668410965 | -3.45290593183483 | 0.00055458236046642 | 0.0494225448269321 |
| LOC1278211 | AGAP007753-PA. Carbohydrate (sugar) transporter. | 2342.8516699155 | 0.419379594208319 | 0.121805514470396 | 3.44302633613723 | 0.00057524329233166 | 0.0497020037793231 |
| LOC1279767 | AGAP003317-PA. | 399.364738127458 | 0.440454008475609 | 0.12803218611525 | 3.44018189363046 | 0.00058132335639957 | 0.0498016763546038 |

**Immune connection: 29/118 (24.6%)**

**JNK has suppressive effect; 29/118 (24.6%)**

**Over represented in hemocytes: 22/118 (18.6%)**

**Total: 51/118 (43.2%)**

**Unique to this data set**

**Shared with IBF only comparison**

**29/118 shared with IBF only comparison, 89 unique to this dataset**

| Gene | Forward | Reverse |
|---|---|---|
| ***RpL19 (AGAP004422)*** | CCAACTCGCGACAAAACATTC | ACCGGCTTCTTGATGATCAGA |
| ***Puc (AGAP004353)*** | GCCTAGTGGTCAGCTGAAGC | GCCTCGATCAGCGACAGACC |
| ***JNK1 (AGAP022950)*** | GCCGAAGAACGACAAACTATGTGC | GCTGTGTTACTGTATCGTATGCG |
| ***TGase2 (AGAP009098)***[a] | CAGCACGACCAAAATGACCTT | TCCTCTAGACAGGTGTCGATTCTTT |
| ***Eater (AGAP012386)*** | AGACTGGTACCGTCAACTGTGG | GAGTTGGTGCAATACCCTGAGC |
| ***Cyp315a1 (AGAP000284)***[b] | CGTCTCGCAACGGCTATCA | CAATCGGATACAGGCGCAAC |

**Supplementary Table 3. qRT primers used in this study.** All primers were used at 300nM except *RpL19* Rev which was used at 900nM [2]. [a] primers reported in [3]. [b] primers reported in [4].

**Supplementary data: Original annotated western blots presented in the manuscript**

**Figure 1a original blots**

Figure 1a original western blot (pJNK)
U=unfed, I=infected blood meal, N=naive blood meal

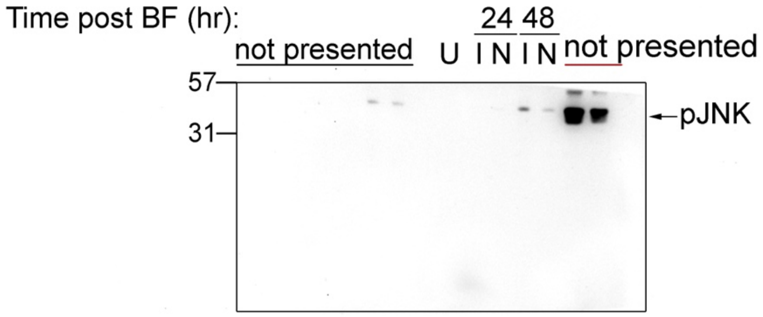


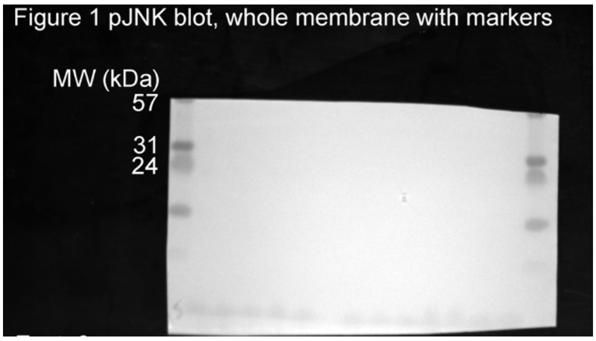


Figure 1a original western blot (pS6K)
U=unfed, I=infected blood meal, N=naive blood meal

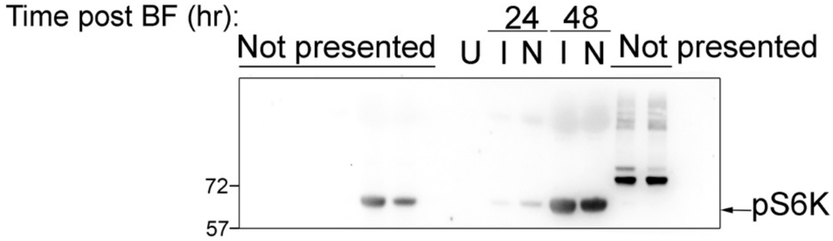


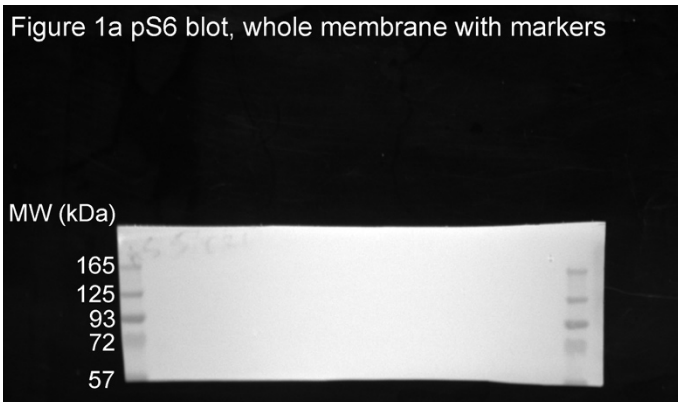

Figure 1a original western blot (actin reprobe)
U=unfed, I=infected blood meal, N=naive blood meal

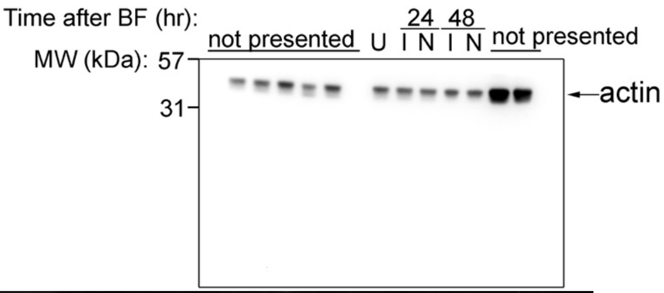


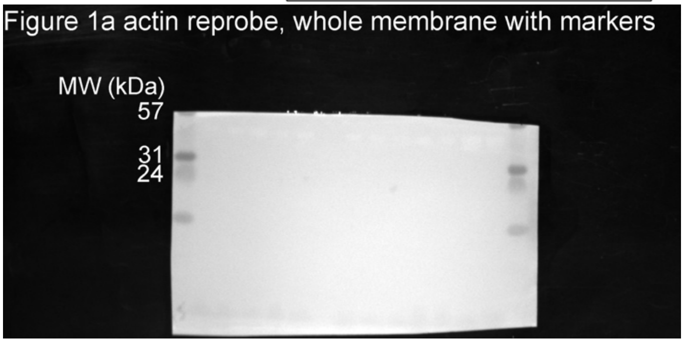

Figure 1a actin reprobe, whole membrane with markers

### Figure 3a original blots

Figure 3a original western blot (pS6K, pJNK, higher exposure)
G=*dsGFP*-injected, P=*dspuc*-injected, P/J=*dspuc/dsJNK*-injected

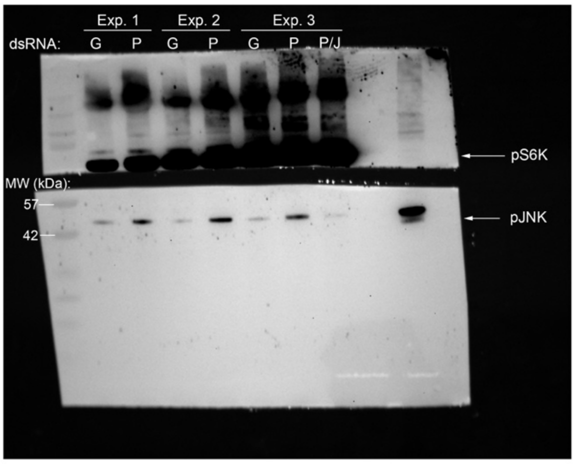


Figure 3a original western blot (pS6K andpJNK blot, low exposure)
G=*dsGFP*-injected, P=*dspuc*-injected, P/J=*dspuc/dsJNK*-injected

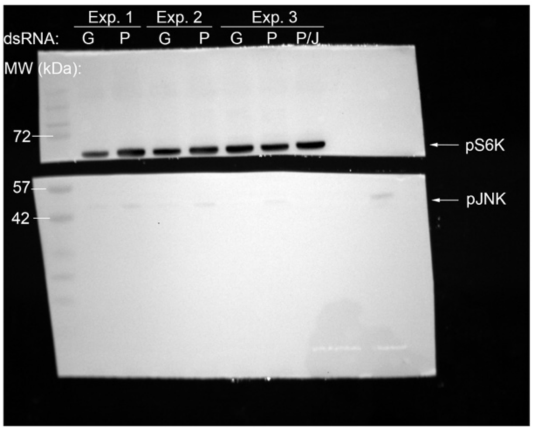


Figure 3a original western blot (Actin reprobe)
G=*dsGFP*-injected, P=*dspuc*-injected, P/J=*dspuc/dsJNK*-injected

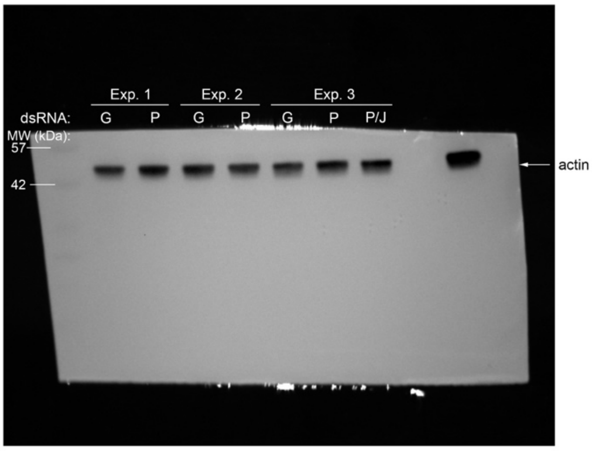

#### Figure 4a original blots

Figure 4a original western blot pJNK vs pS6K
Ambient temperature and time after blood feeding indicated

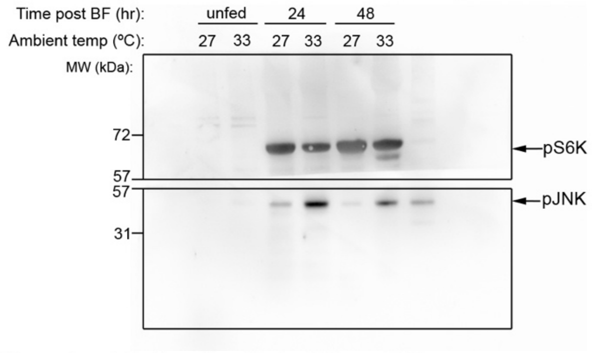


Figure 4a original western blot pJNK vs pS6K.
Whole membrane with markers

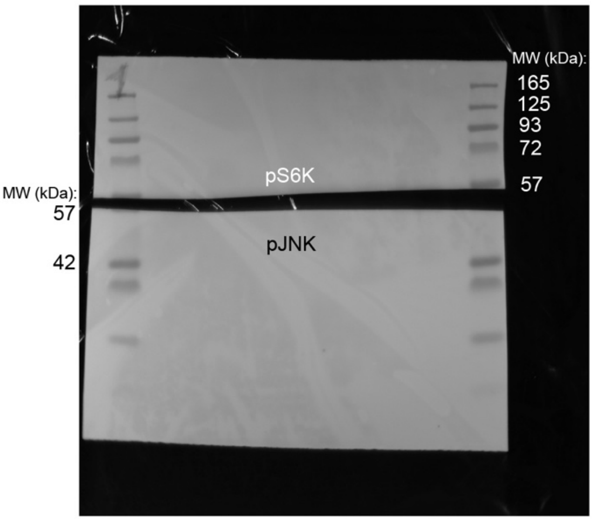


Figure 4a original western blot (actin reprobe)
Ambient temperature and time after blood feeding indicated

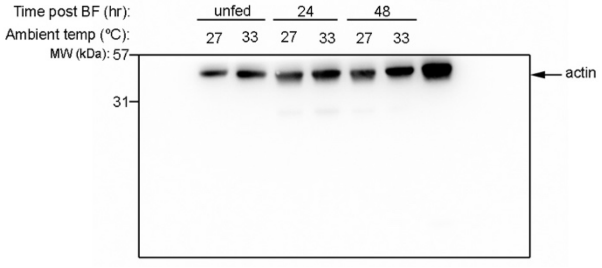

#### Supplementary Figure 6a

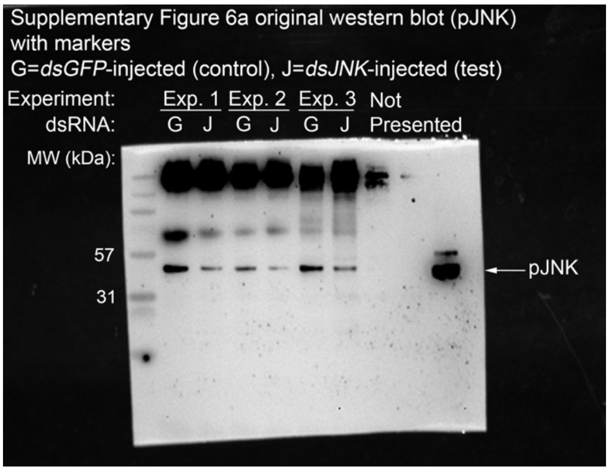


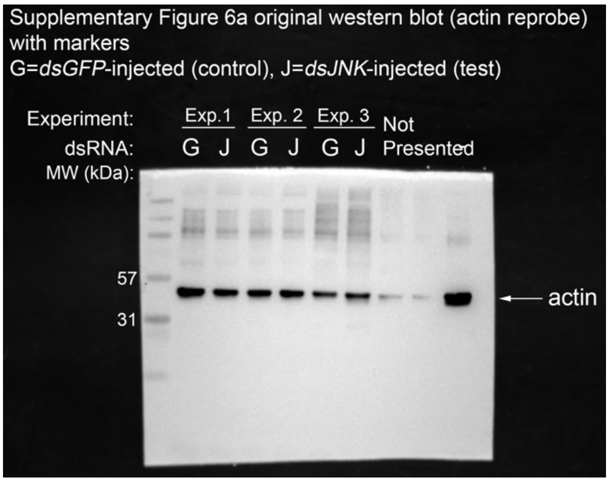
